# An Autoantigen-ome from HS-Sultan B-Lymphoblasts Offers a Molecular Map for Investigating Autoimmune Sequelae of COVID-19

**DOI:** 10.1101/2021.04.05.438500

**Authors:** Julia Y. Wang, Wei Zhang, Victor B. Roehrl, Michael W. Roehrl, Michael H. Roehrl

## Abstract

To understand how COVID-19 may induce autoimmune diseases, we have been compiling an atlas of COVID-autoantigens (autoAgs). Using dermatan sulfate (DS) affinity enrichment of autoantigenic proteins extracted from HS-Sultan lymphoblasts, we identified 362 DS-affinity proteins, of which at least 201 (56%) are confirmed autoAgs. Comparison with available multi-omic COVID data shows that 315 (87%) of the 362 proteins are affected in SARS-CoV-2 infection via altered expression, interaction with viral components, or modification by phosphorylation or ubiquitination, at least 186 (59%) of which are known autoAgs. These proteins are associated with gene expression, mRNA processing, mRNA splicing, translation, protein folding, vesicles, and chromosome organization. Numerous nuclear autoAgs were identified, including both classical ANAs and ENAs of systemic autoimmune diseases and unique autoAgs involved in the DNA replication fork, mitotic cell cycle, or telomerase maintenance. We also identified many uncommon autoAgs involved in nucleic acid and peptide biosynthesis and nucleocytoplasmic transport, such as aminoacyl-tRNA synthetases. In addition, this study found autoAgs that potentially interact with multiple SARS-CoV-2 Nsp and Orf components, including CCT/TriC chaperonin, insulin degrading enzyme, platelet-activating factor acetylhydrolase, and the ezrin-moesin-radixin family. Furthermore, B-cell-specific IgM-associated ER complex (including MBZ1, BiP, heat shock proteins, and protein disulfide-isomerases) is enriched by DS-affinity and up-regulated in B-cells of COVID-19 patients, and a similar IgH-associated ER complex was also identified in autoreactive pre-B1 cells in our previous study, which suggests a role of autoreactive B1 cells in COVID-19 that merits further investigation. In summary, this study demonstrates that virally infected cells are characterized by alterations of proteins with propensity to become autoAgs, thereby providing a possible explanation for infection-induced autoimmunity. The COVID autoantigen-ome provides a valuable molecular resource and map for investigation of COVID-related autoimmune sequelae and considerations for vaccine design.

## Introduction

The novel coronavirus SARS-CoV-2 has caused the worldwide COVID-19 pandemic with hundreds of millions infected and high morbidity and mortality. A significant proportion of patients who have recovered from the acute viral infection of COVID-19 continue to suffer from lingering health problems, so called “long COVID” syndrome. It is yet unknown how long the COVID aftereffects will persist, and more importantly, what the underlying causative mechanisms of long COVID syndrome are. The acute phase of COVID-19 is accompanied by various autoimmune responses, and autoimmune diseases, which tend to be chronic and debilitating, are major concerns of COVID-19 sequelae. To understand how SARS-CoV-2 infection may induce autoimmunity and how diverse the autoimmune disorders could be, we have started to compile a comprehensive atlas of COVID autoantigens (autoAgs) and autoantibodies (autoAbs), the root elements of autoimmunity [1, 2].

We have developed a unique DS-affinity enrichment strategy for autoAg discovery [1–8]. We discovered that dermatan sulfate (DS), a glycosaminoglycan that is abundant in skin and soft connective tissues and that is involved in wound healing and tissue repair, has affinity to autoAgs [3, 4]. Because of this affinity, DS binds autoAgs to form non-covalent DS-autoAg complexes, which transforms non-antigenic singular self-molecules into antigenic non-self-complexes [3]. DS-autoAg complexes are capable of engaging autoreactive B-cell receptors (autoBCRs) via a two-step process: (i) DS-autoAg complexes bind autoBCRs on autoreactive B1-cells via autoAg-autoBCR specificity; (ii) DS enters cells by (DS-autoAg)-autoBCR complex internalization and recruits a cascade of molecules to stimulate autoreactive B1-cells [3, 5]. In particular, DS recruits GTF2I that is required for IGH gene expression and IgH-associated multiprotein complexes in the endoplasmic reticulum (ER) to facilitate autoAb production [5]. Therefore, any self-molecule with DS-affinity has a propensity to be transformed by DS into an autoantigenic DS-autoAg complex [5]. Based on this unifying principle of DS-autoAg affinity, we have discovered and catalogued known and putative autoAgs from different cells and organs [1, 2, 6–8].

Autoantibodies, which target autoAgs, have been found in a significant portion of COVID-19 patients. In a cohort study of 147 hospitalized COVID-19 patients, autoAbs are detected in ∼50% of the patients, and antinuclear autoAbs are detected in ∼25% of patients, with the target autoAgs associated with myositis, systemic sclerosis, and connective tissue disease overlap syndromes [9]. In another study of 987 COVID- 19 patients with life-threatening pneumonia, over 10% developed autoAbs against interferons, which likely neutralized their ability to block SARS-CoV-2 infection [10]. Although COVID-19 is typically mild or asymptomatic in children, multisystem inflammatory syndrome and multiple autoAbs developed in a portion of infected children [11, 12]. In a study of COVID-19 patients with unexplained neurological symptoms, anti-neuronal autoAbs were detected in sera or cerebrospinal fluid of all patients [13]. Antinuclear autoAbs, the most frequently tested autoAbs in clinical screening for autoimmune diseases such as lupus, Sjögren syndrome, scleroderma, or polymyositis, are found in 20-50% of COVID-19 patients [14–16].

Viral infections have long been regarded as culprits of autoimmune diseases. However, it has remained unclear how infections induce autoimmune diseases. In this study, we investigated HS-Sultan cells, a B-cell lymphoblast line isolated from Burkitt’s lymphoma of a 7-year-old boy. HS-Sultan cells are infected with and immortalized by Epstein-Barr virus (EBV) and carry the viral DNA sequence. From the proteome extracts of HS-Sultan cells, we identified a putative DS-affinity autoantigen-ome of 362 proteins, of which 201 are confirmed autoAgs by means of corresponding specific autoAbs reported in the literature. By comparing this autoantigen-ome with proteins affected by SARS-CoV-2 infection derived from multi-omic studies compiled in Coronascape [17–38], we identified 315 DS-affinity proteins and 186 confirmed autoAgs. When host cells are infected, numerous molecules undergo significant changes via altered expression, modification, or degradation. When the infected cells die, these altered molecules are released, and those with DS-affinity may become associated with DS and transform into immunogenic autoAg-DS complexes. This study illustrates that viral infections can profoundly change the host cell autoantigen-ome, result in a large repertoire of potential autoAgs, and may consequently lead to autoimmune disease.

## Results and Discussion

### DS-affinity autoantigen-ome of HS-Sultan cells

HS-Sultan cells were cultured, harvested, and lysed. Total proteins were extracted from lysates and fractionated on DS-Sepharose affinity resins. Proteins with no or weak DS-affinity were removed from the resins with 0.2 M NaCl, and those with intermediate to strong DS-affinity were eluted first with 0.5 M and then with 1.0 M NaCl. Proteins in the DS-affinity fractions were collected, desalted, digested, and sequenced by mass spectrometry. A total of 362 proteins were identified, with the majority present in the 0.5 M NaCl elution. Proteins that eluted with 1.0 M NaCl possess very strong DS-affinity and include some of the classical nuclear autoAgs, e.g., histones, TOP1, Sm-D3, and 60S acidic ribosomal protein P0. Other proteins with strong DS-affinity include both known autoAgs (vimentin, ATP synthetase ATP5B, and PABPC1) and unknown ones (RPL10A, L15, RPS27A, and mitochondrial single-stranded DNA binding protein SSBP1).

Of the 362 DS-affinity proteins identified from HS-Sultan cells, 201 (55.5%) are confirmed humoral autoAgs based on prior literature reports of specific autoantibodies (see references in Table 1). These autoAgs and their respective autoantibodies are found in a wide spectrum of autoimmune diseases and cancer. The number of actual autoAgs is likely much greater, as most of the unconfirmed proteins have structural resemblance to known autoAgs. For example, SSBP1 is structurally and functionally similar to the classical lupus autoAg SSB, but it has not formally been identified as an autoAg. As another example, nucleosome assembly protein 1-like 1 and 4 (NAP1L1, NAP1L4) are identified by DS-affinity but unconfirmed as autoAgs, whereas their close relative NAP1L3 has been reported as an autoAg. Due to the structural similarity of many DS-affinity proteins with known autoAgs, it is likely that there are additional yet-to-be discovered (considered putative) autoAgs in this group.

**Table 1.**
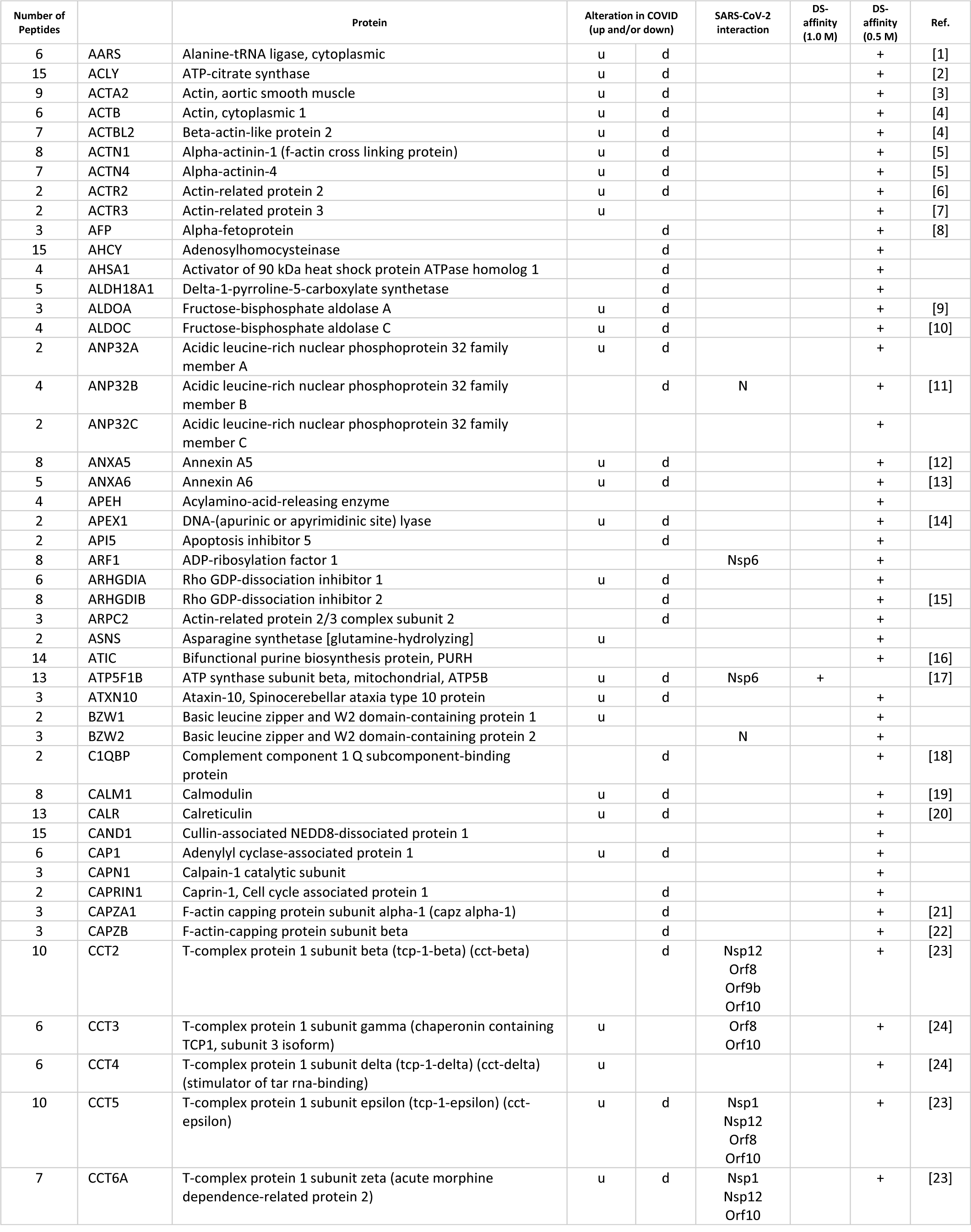
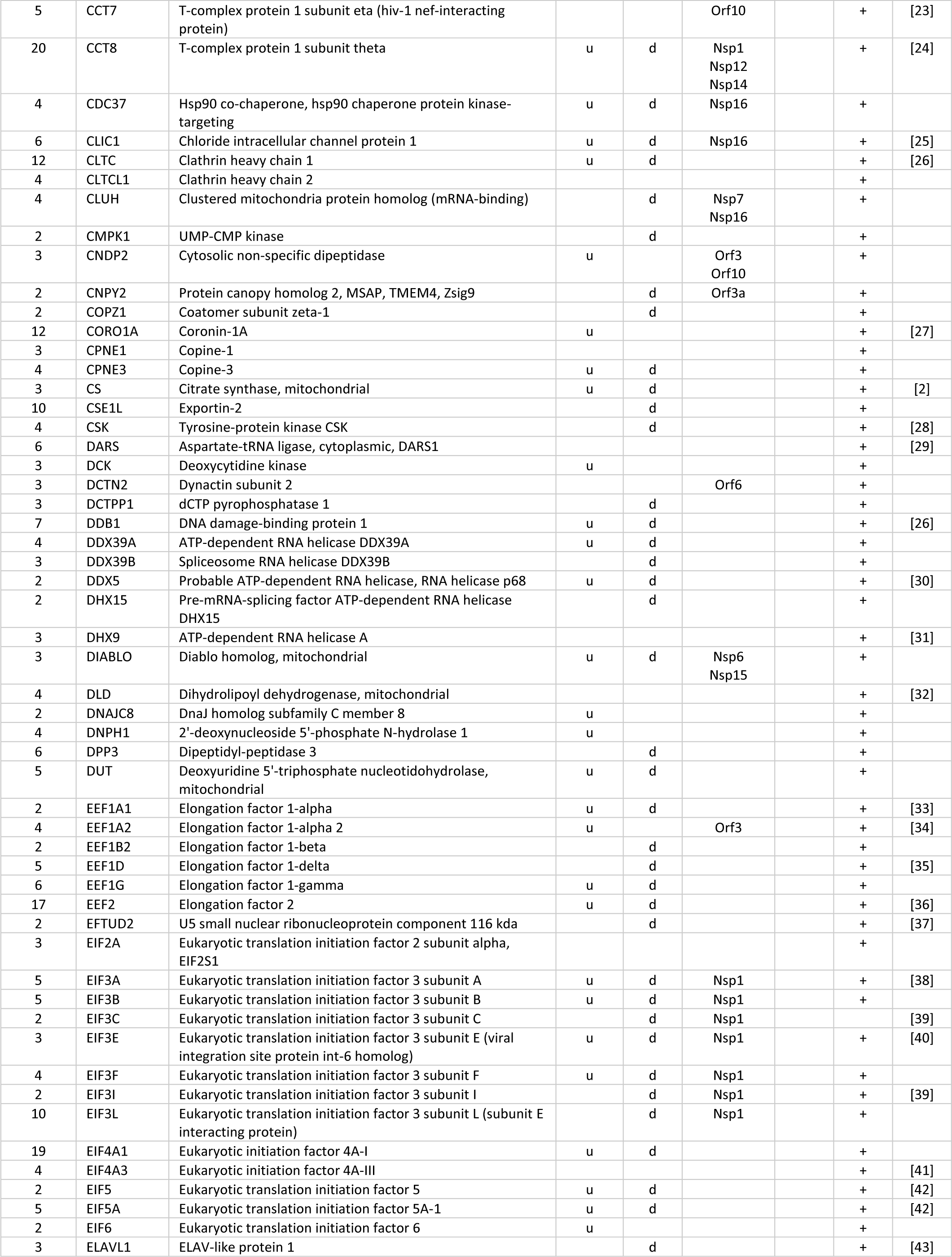
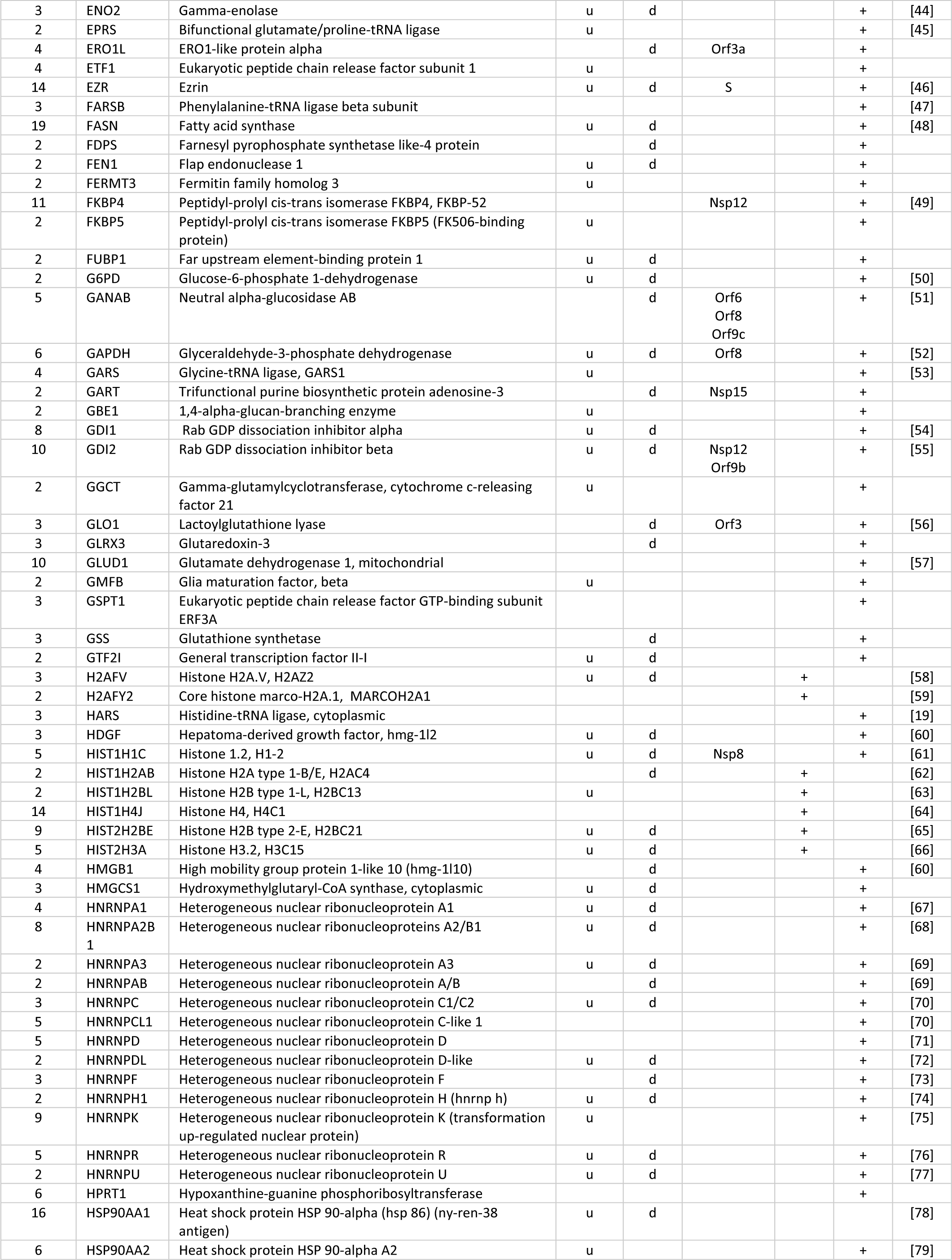
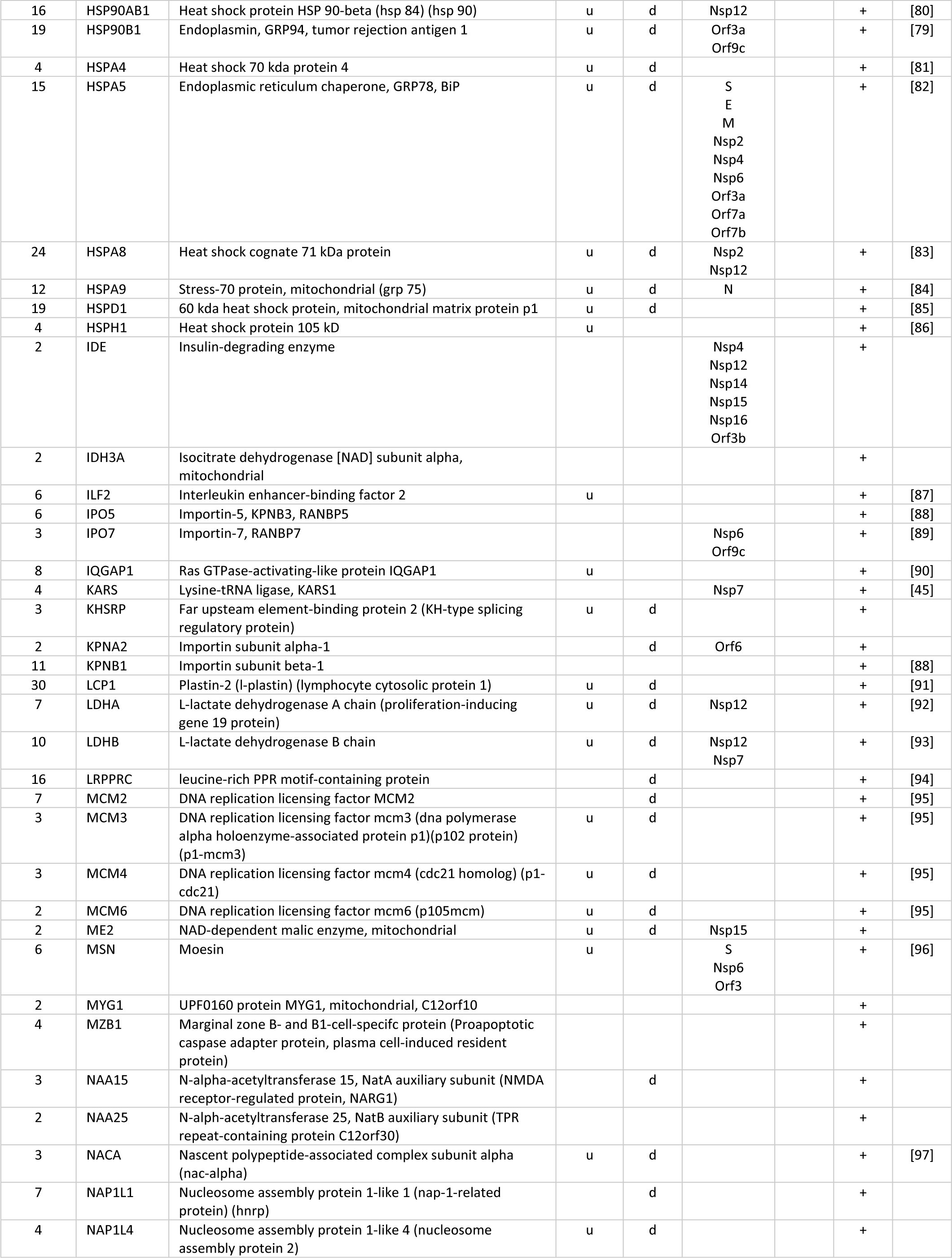
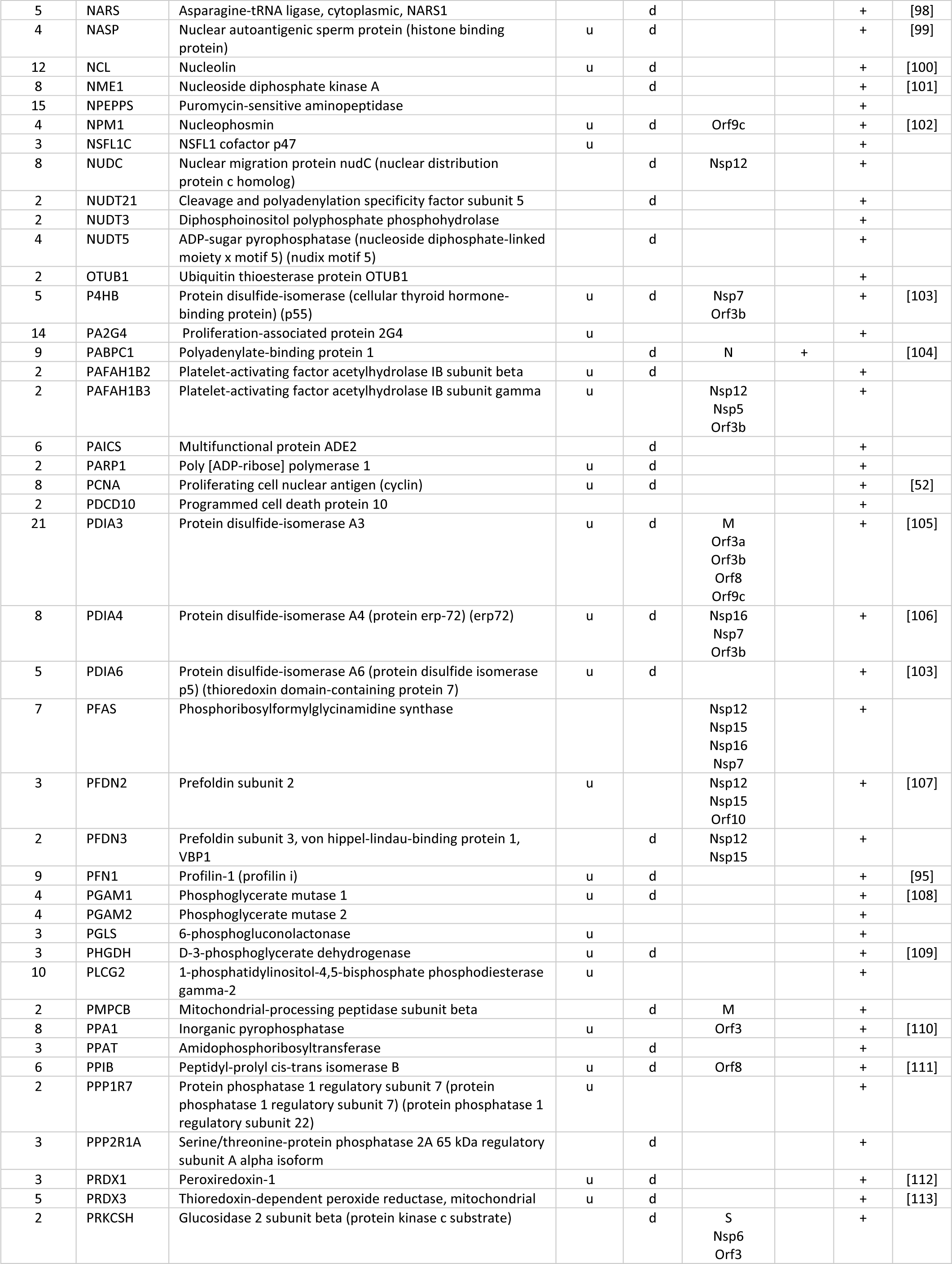
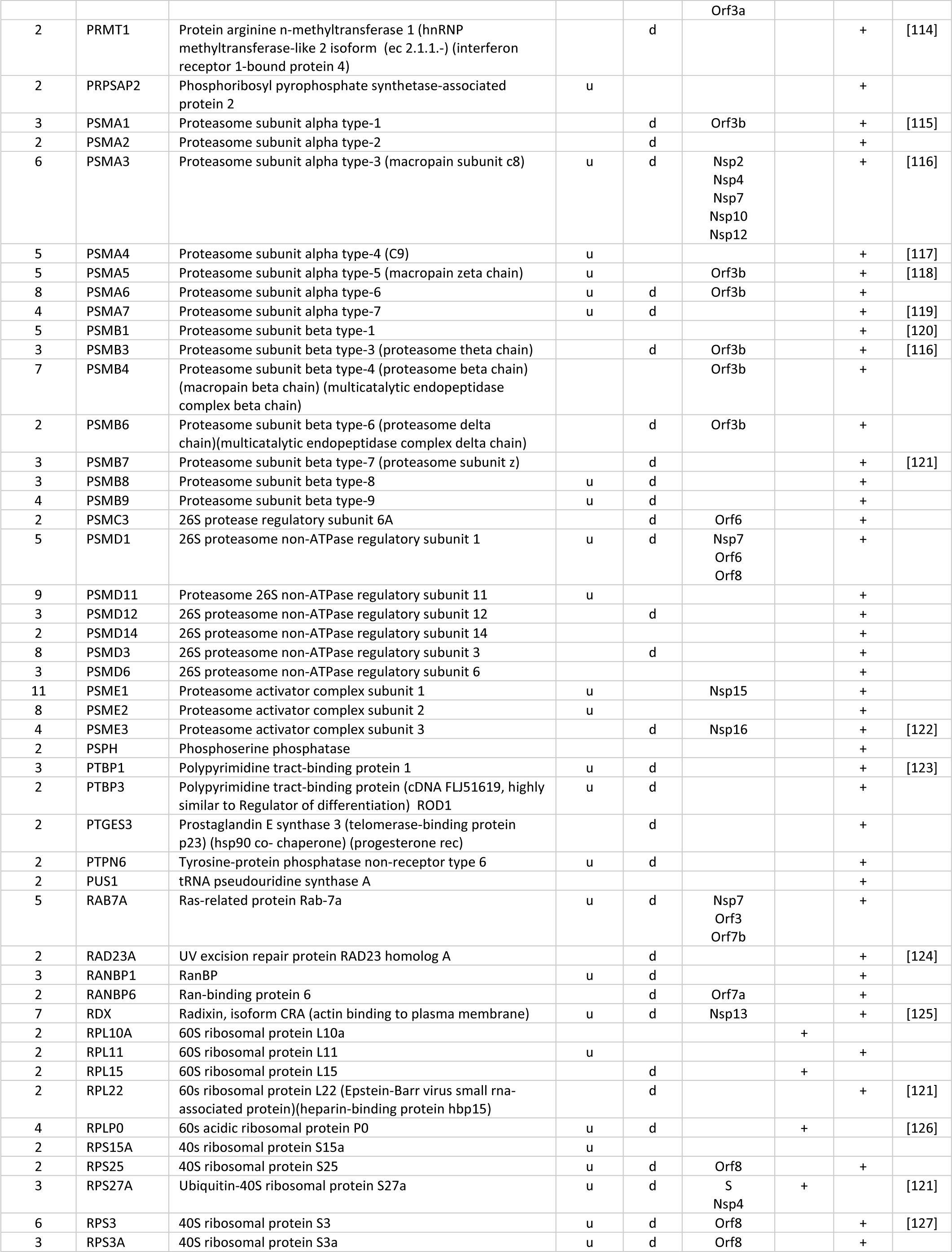
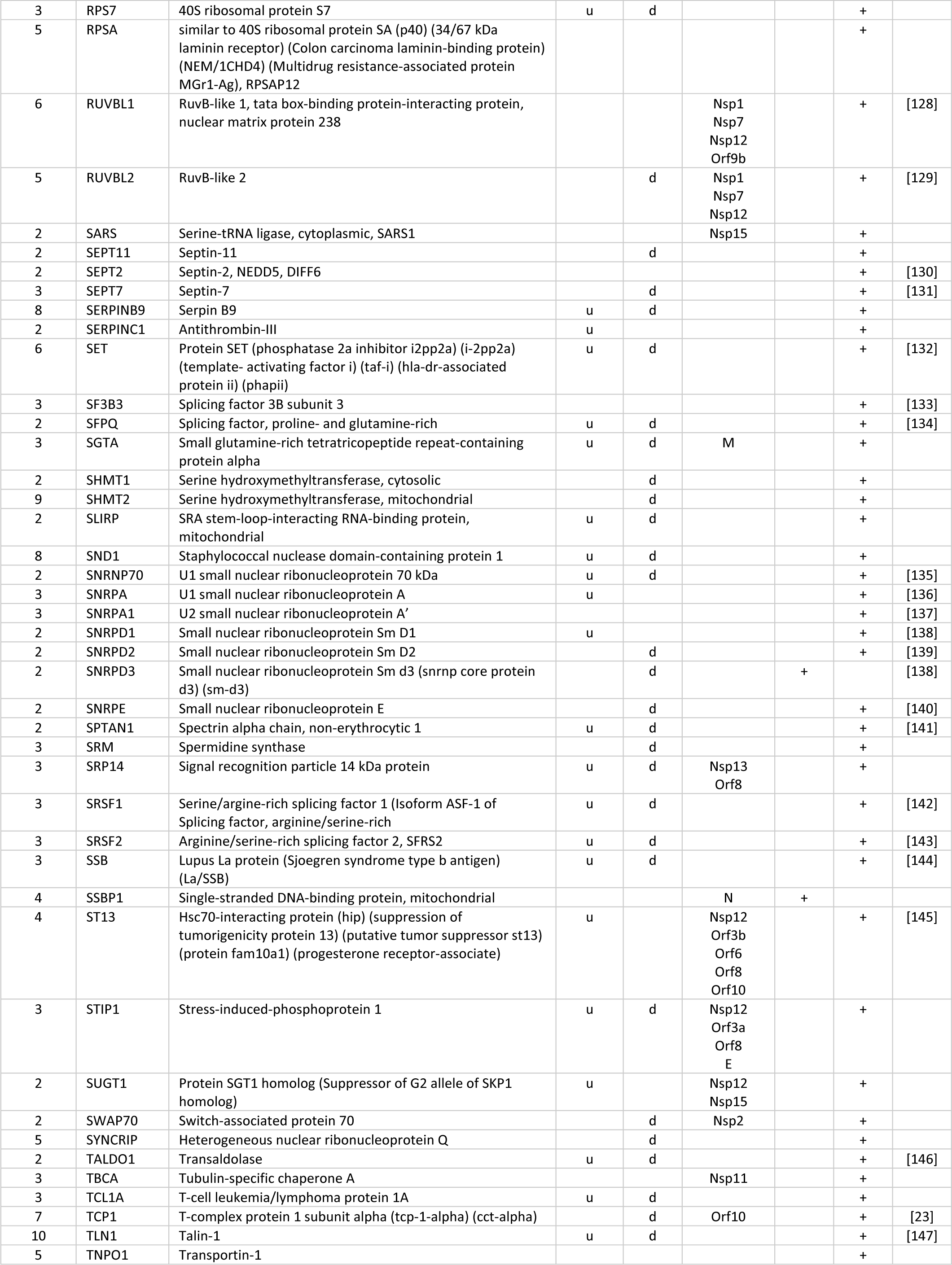
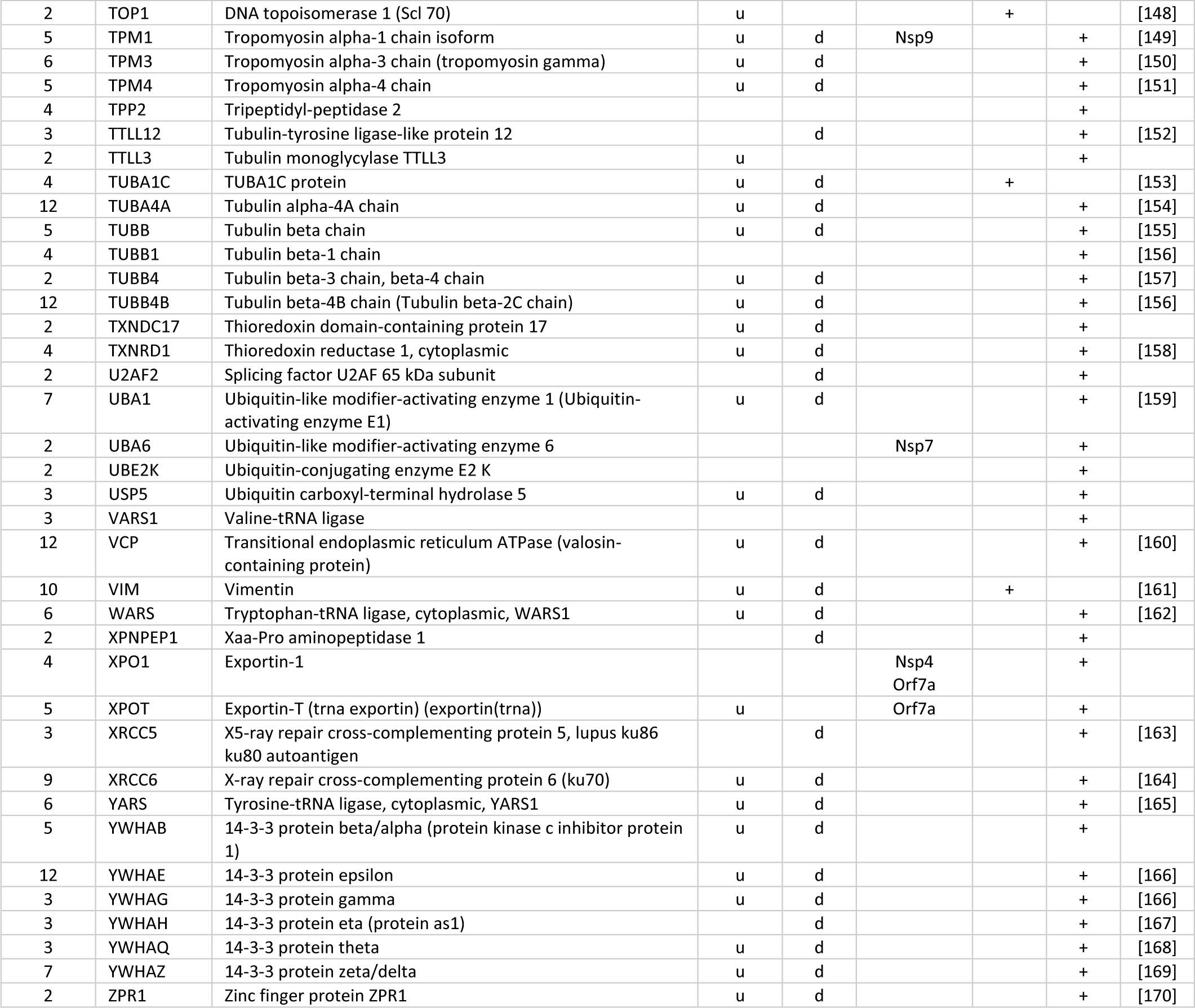
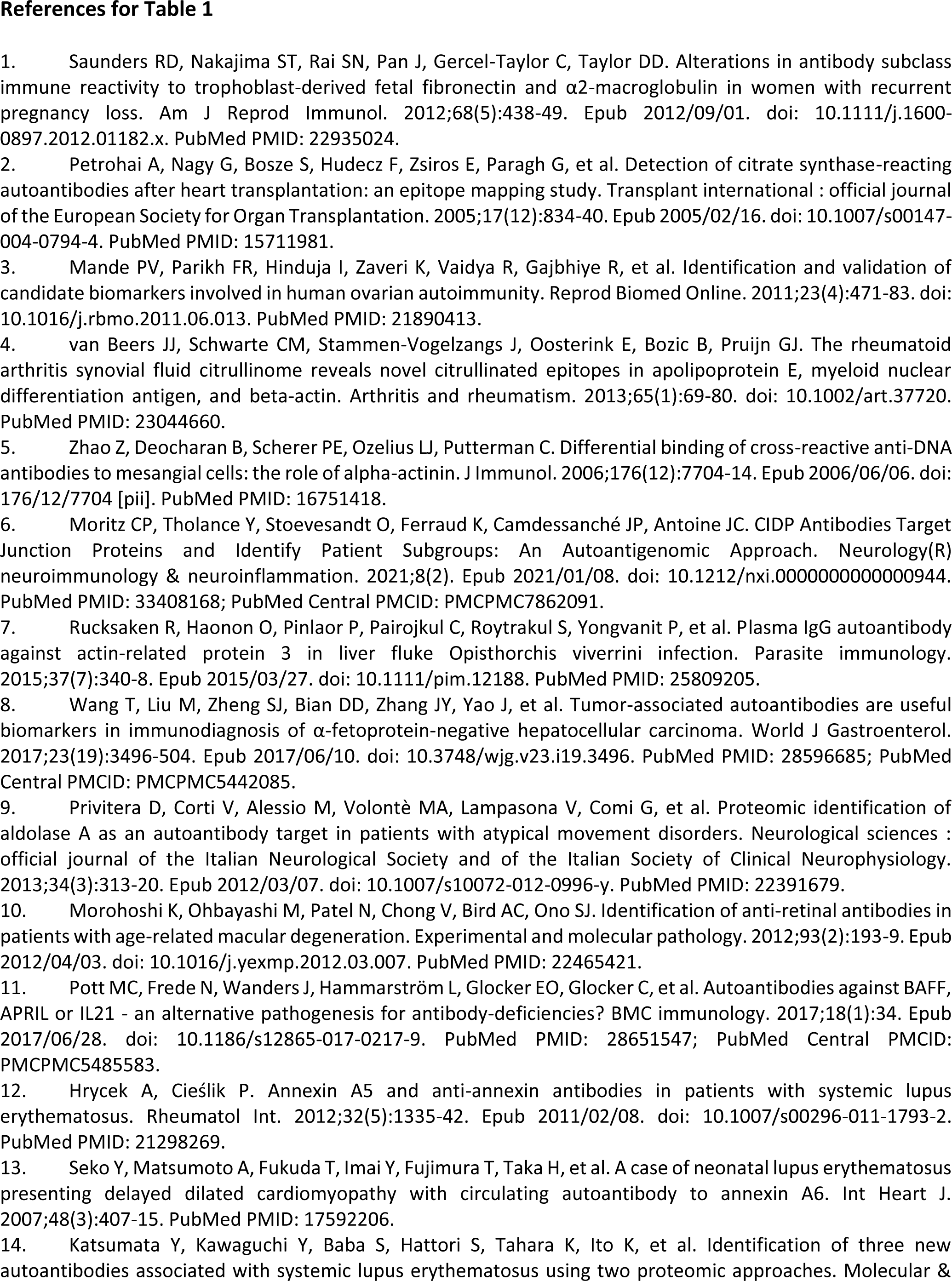
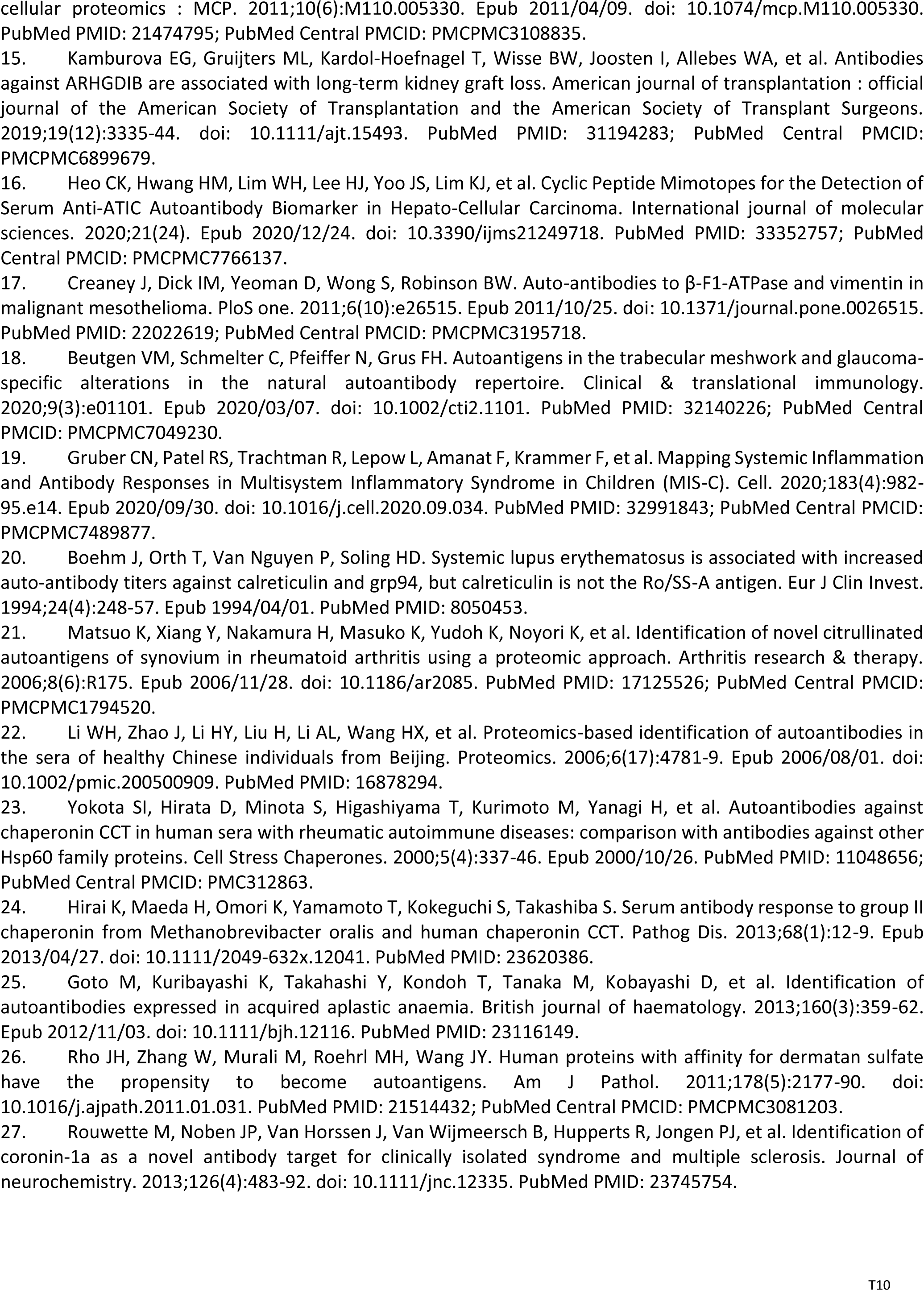
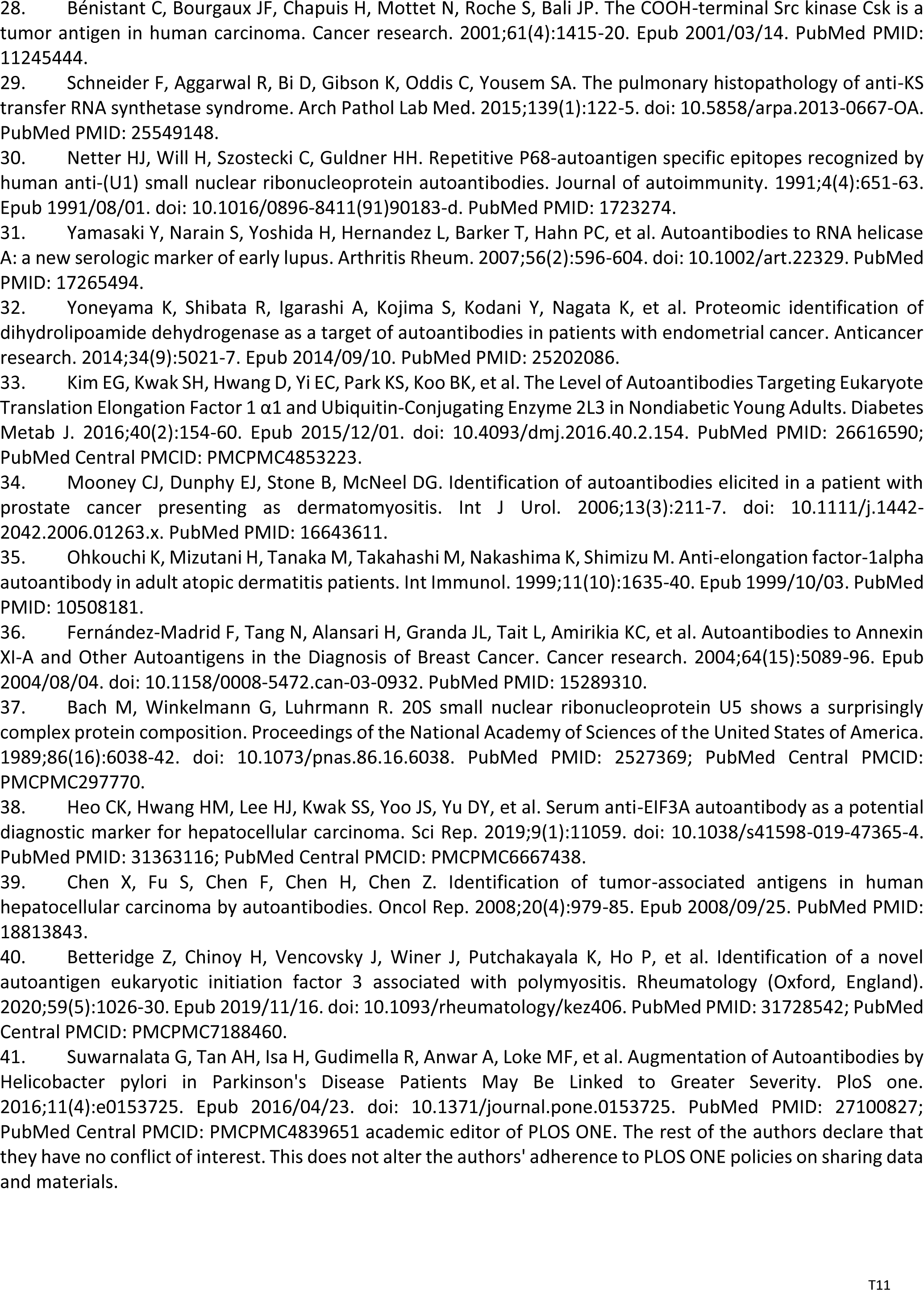
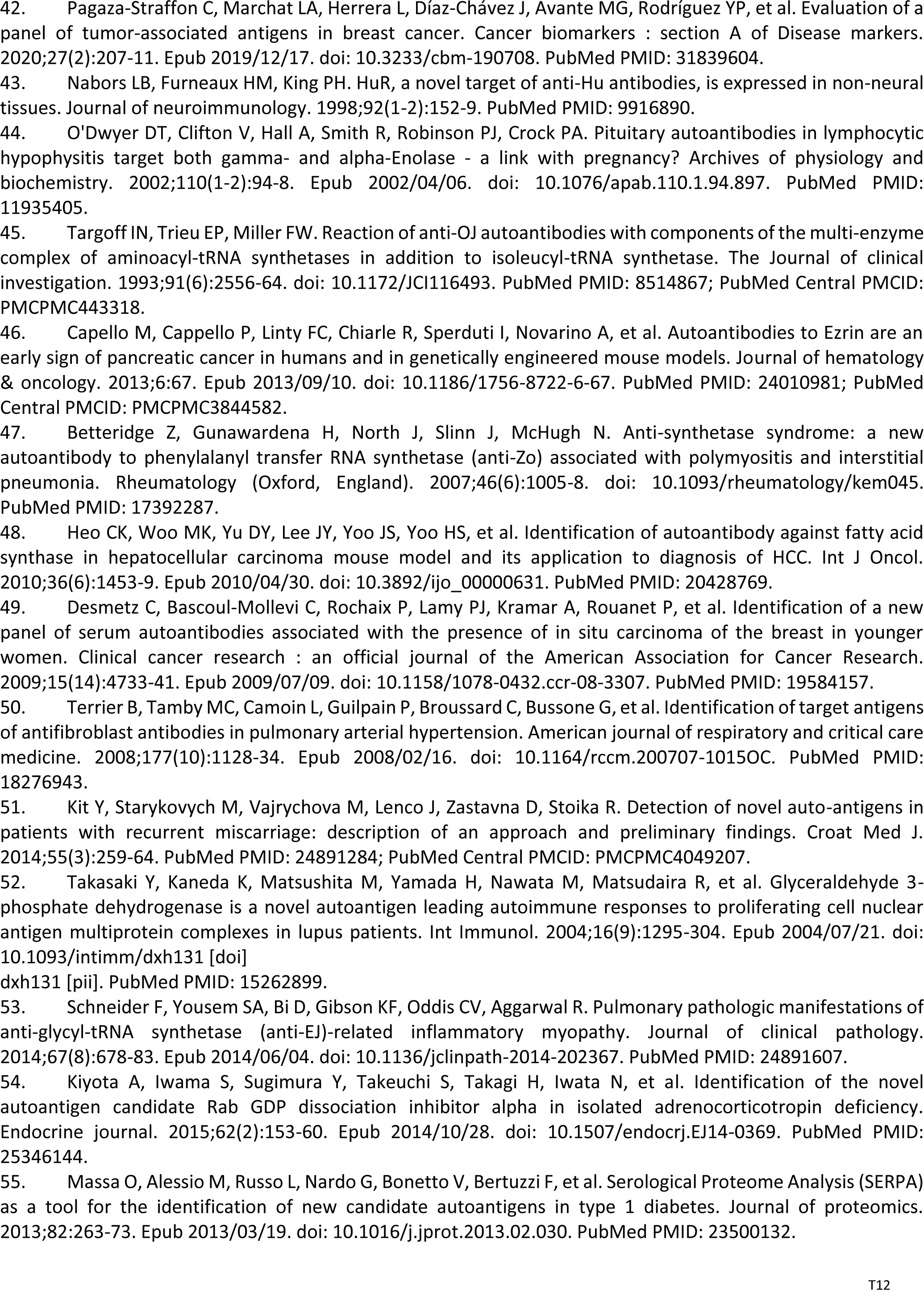
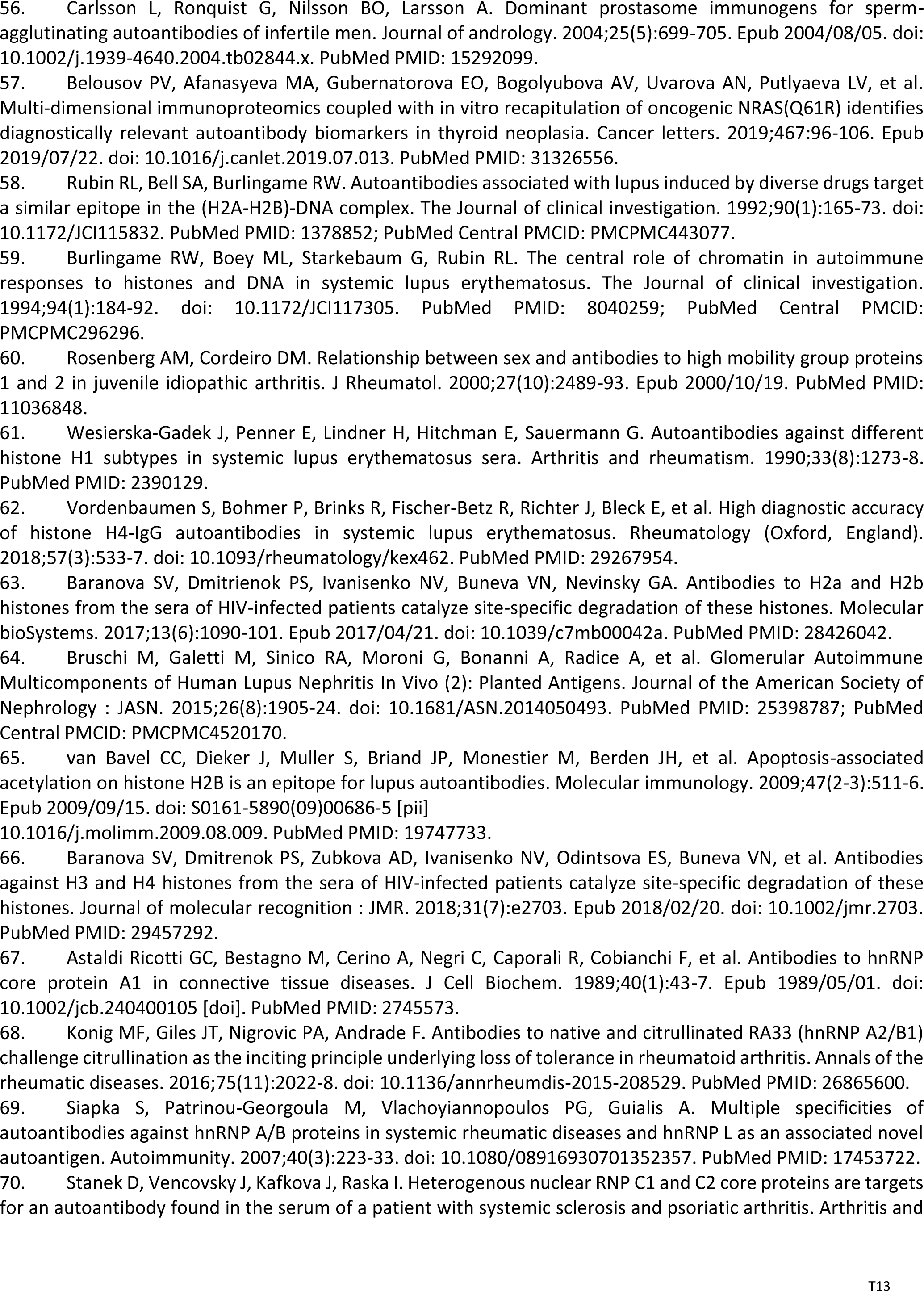
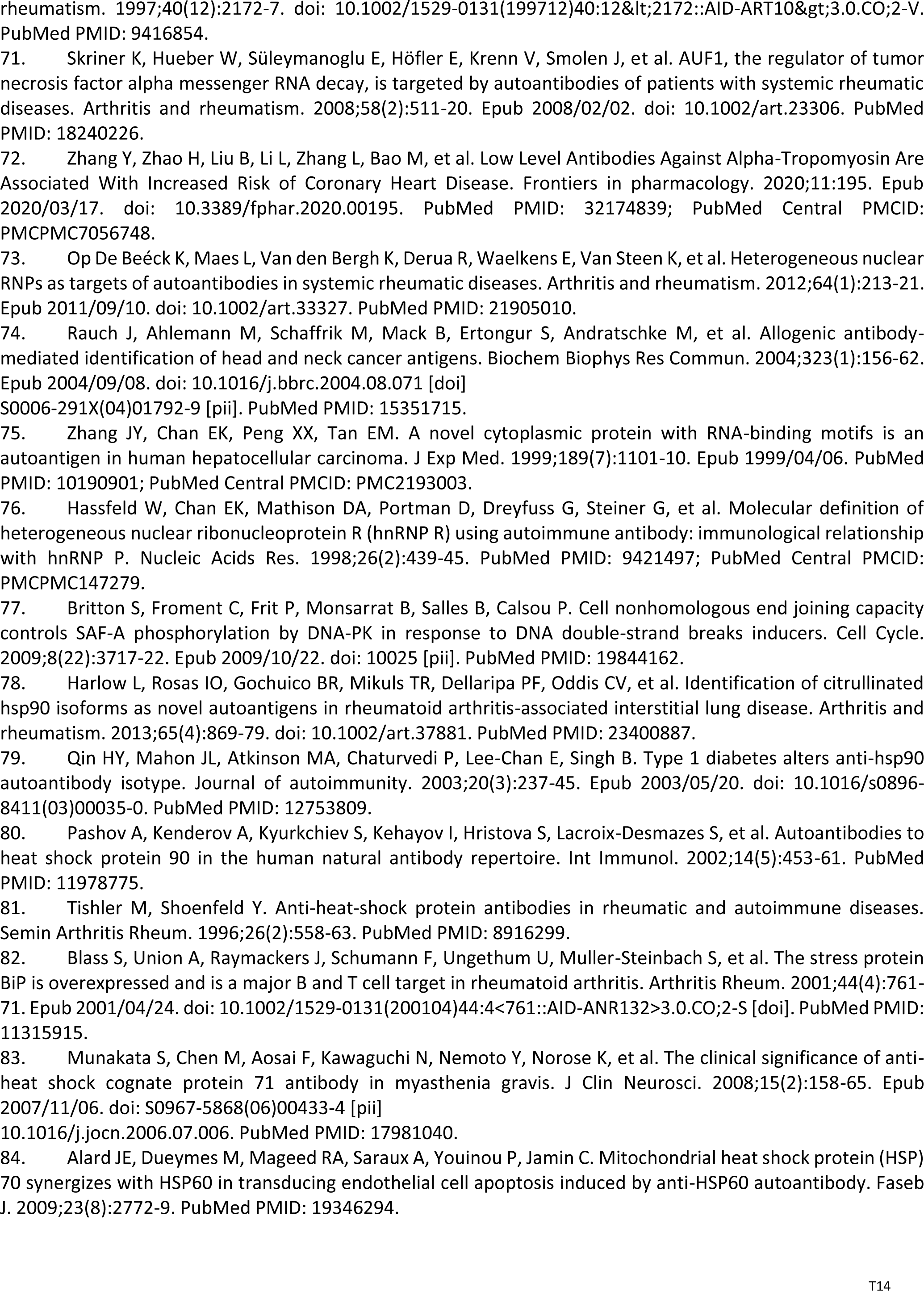
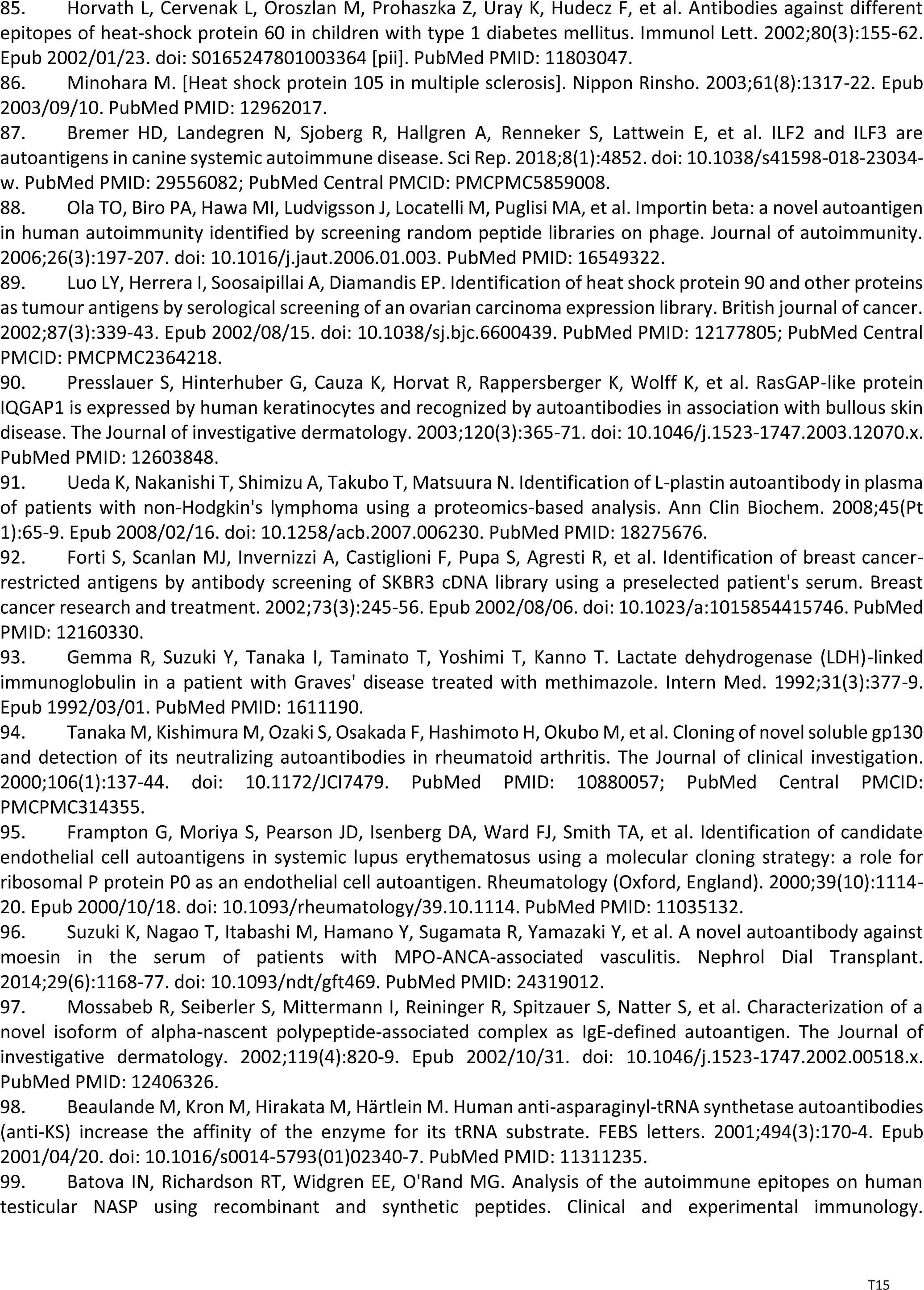
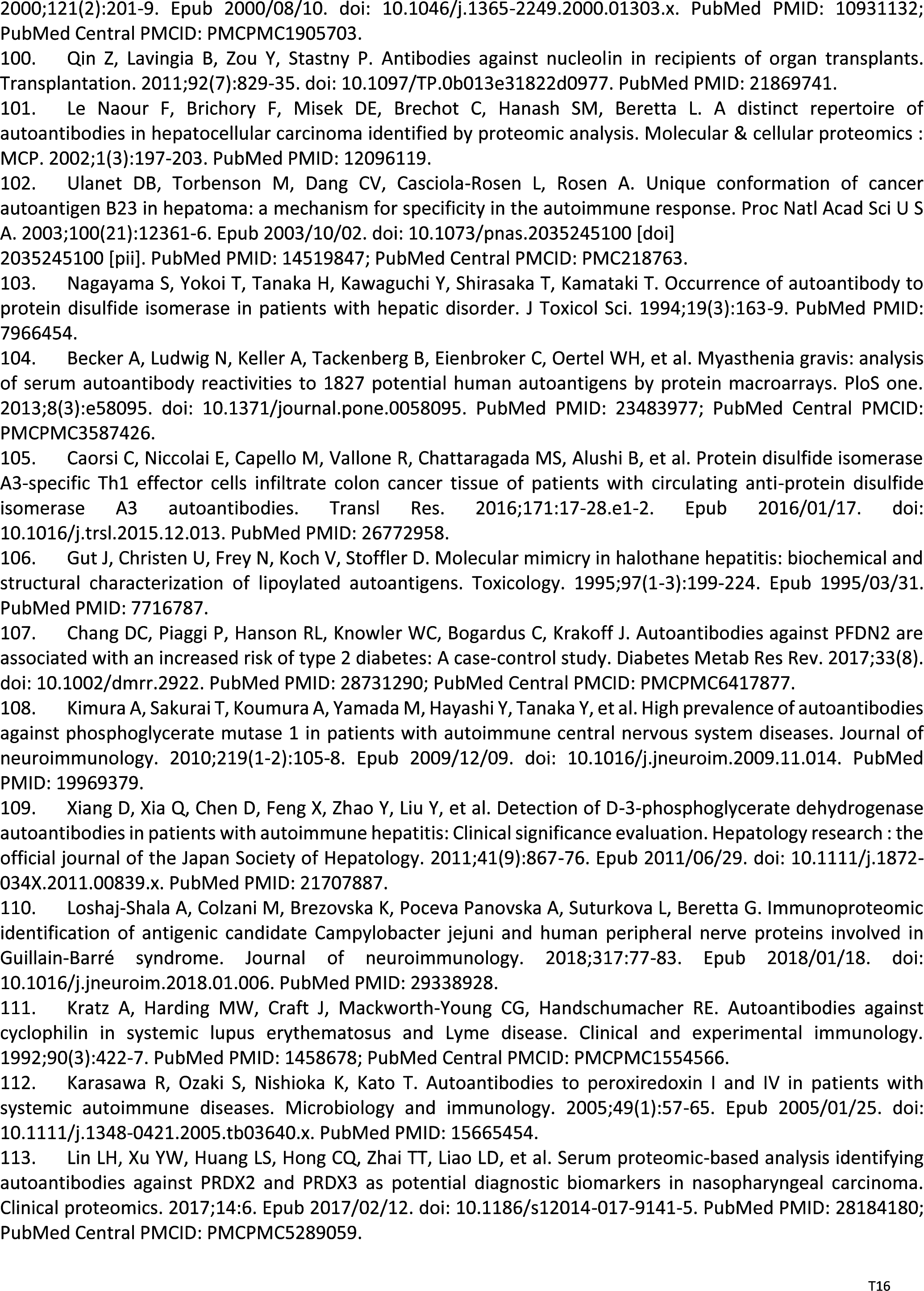
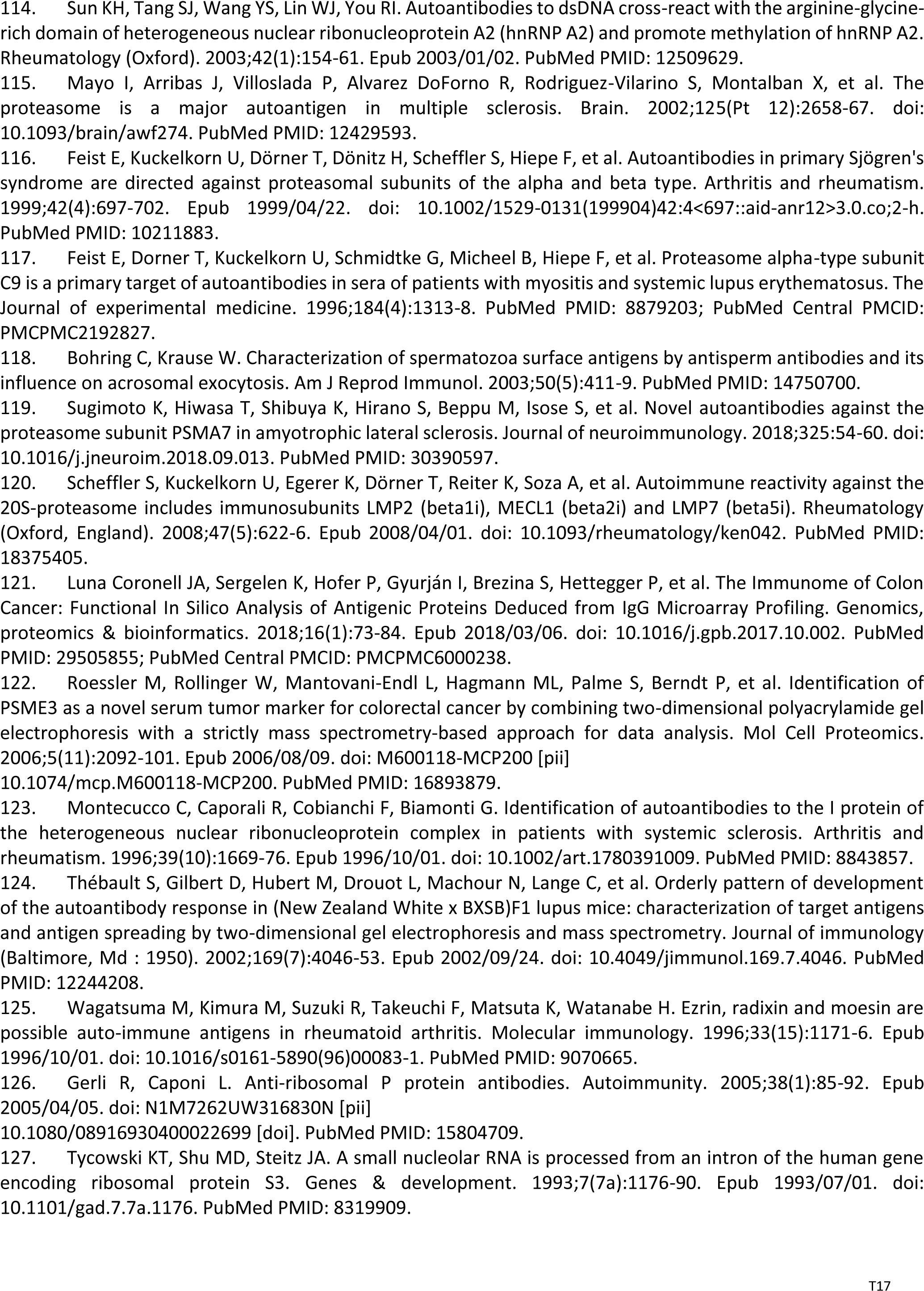
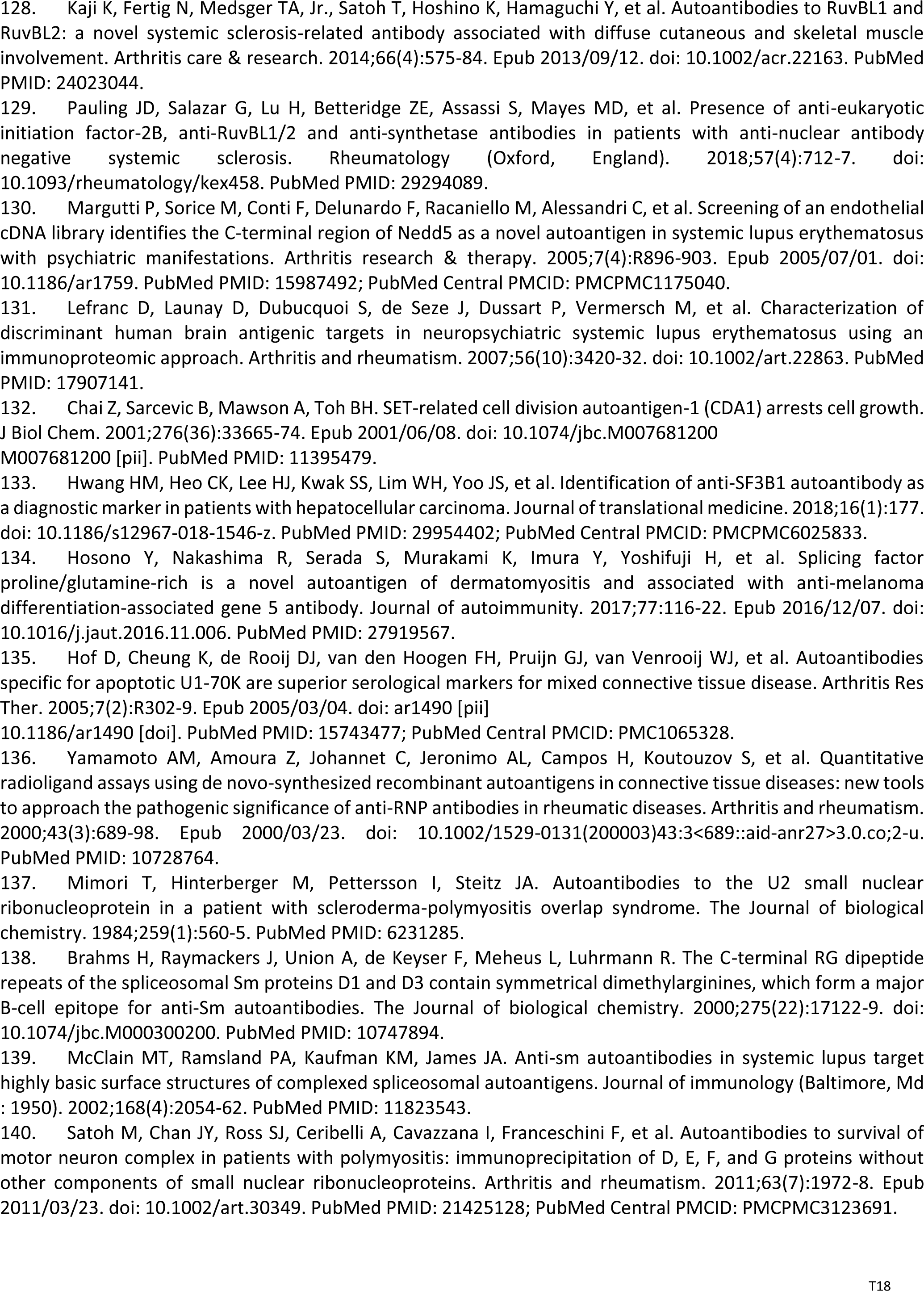
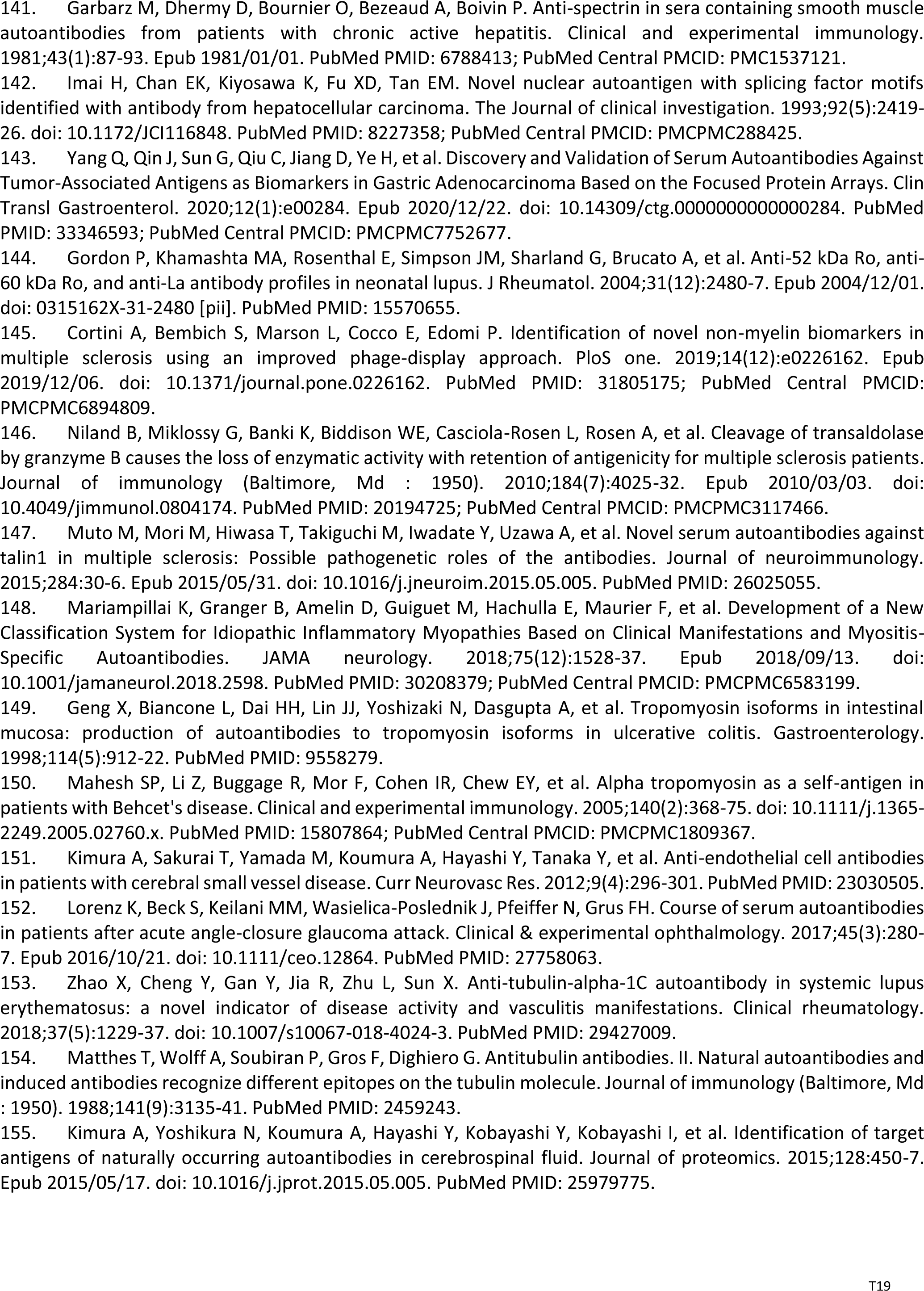
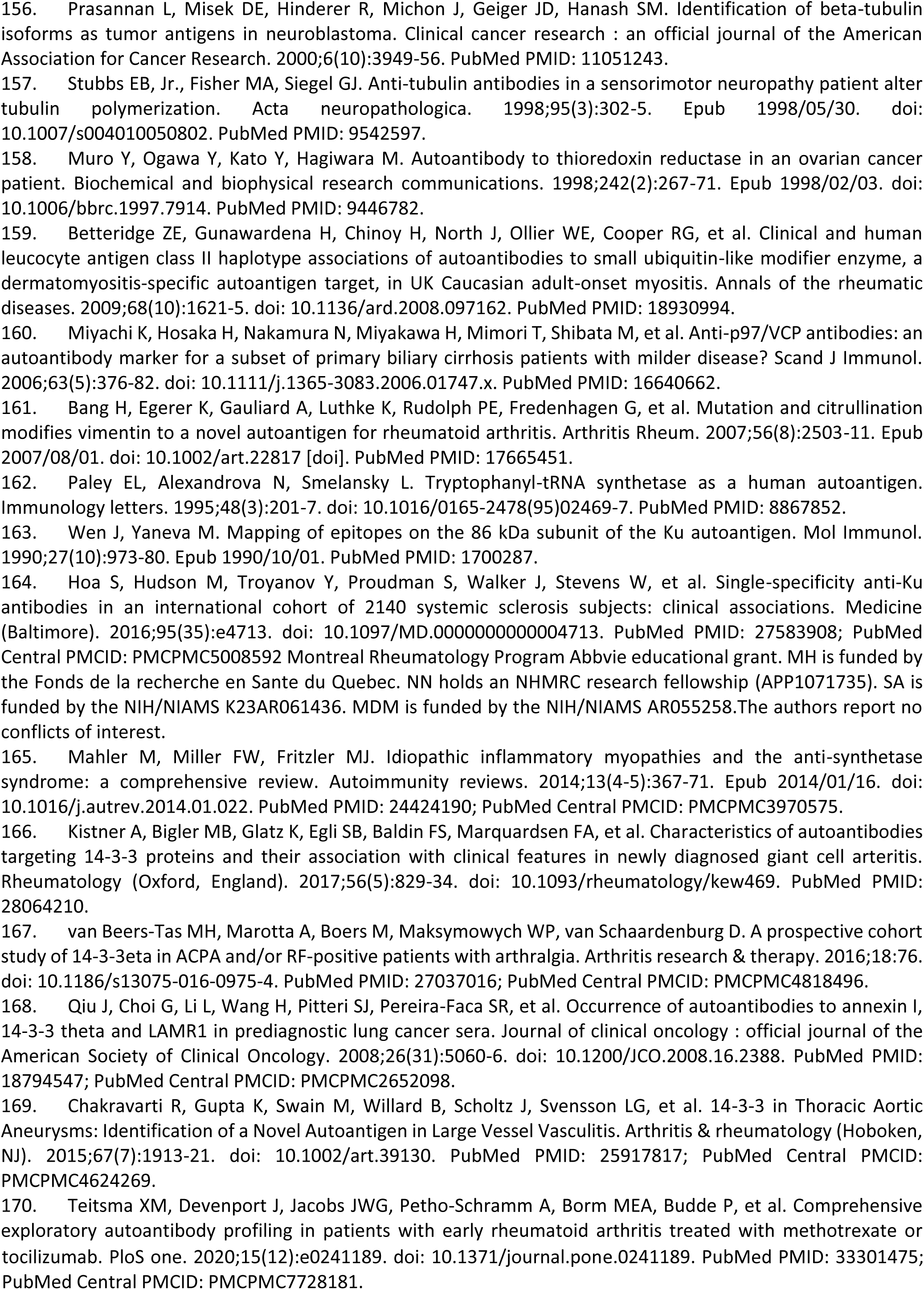
DS-affinity autoantigens from HS-Sultan cells and their alteration in COVID-19

The DS-affinity autoantigen-ome, which includes confirmed and putative autoAgs, is not a random collection of proteins but functionally highly connected. As shown by protein-protein interaction analysis, the DS-affinity autoantigen-ome possesses significantly more interactions than expected (3,105 interactions at high confidence level vs. 1,249 expected; PPI enrichment p-value <1.0E-16). Based on Gene Ontology (GO) biological process analysis, the DS-affinity autoantigen-ome of HS-Sultan cells is significantly associated with RNA slicing, translation, peptide biosynthesis, protein folding, proteolysis, biosynthesis and metabolism of nucleobase-containing small molecules (e.g., nucleobase, nucleoside, and nucleotide phosphate), cytoskeleton organization, and chromosome organization (Fig. 1). Pathway and process enrichment analysis reveals that it is also significantly associated with neutrophil degranulation, nucleocytoplasmic transport, kinase maturation complex, metabolic reprogramming, and IL-12 mediated signaling (Fig. 2A).

**Fig. 1.**
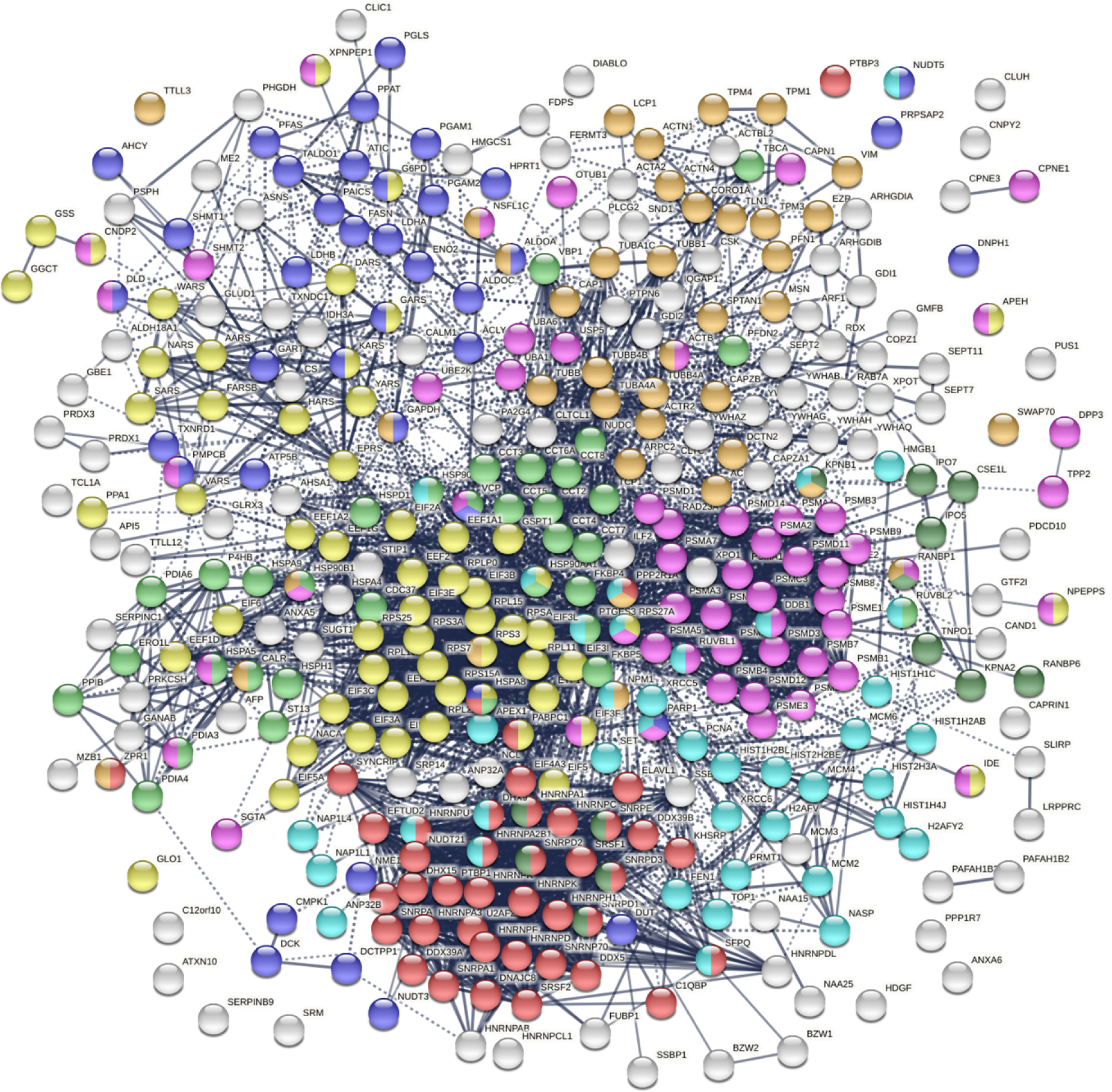
The autoantigen-ome of HS-Sultan cells identified by DS-affinity. Marked proteins are associated with peptide biosynthesis and catabolic process (58 proteins, yellow), protein folding (34 proteins, light green), RNA splicing (41 proteins, red), nucleobase-containing small molecule metabolic process (39 proteins, blue), proteolysis (55 proteins, pink), import into nuclear (13 proteins, dark green), cytoskeleton organization (39 proteins, amber), and chromosome organization (40 proteins, aqua).

**Fig. 2.**
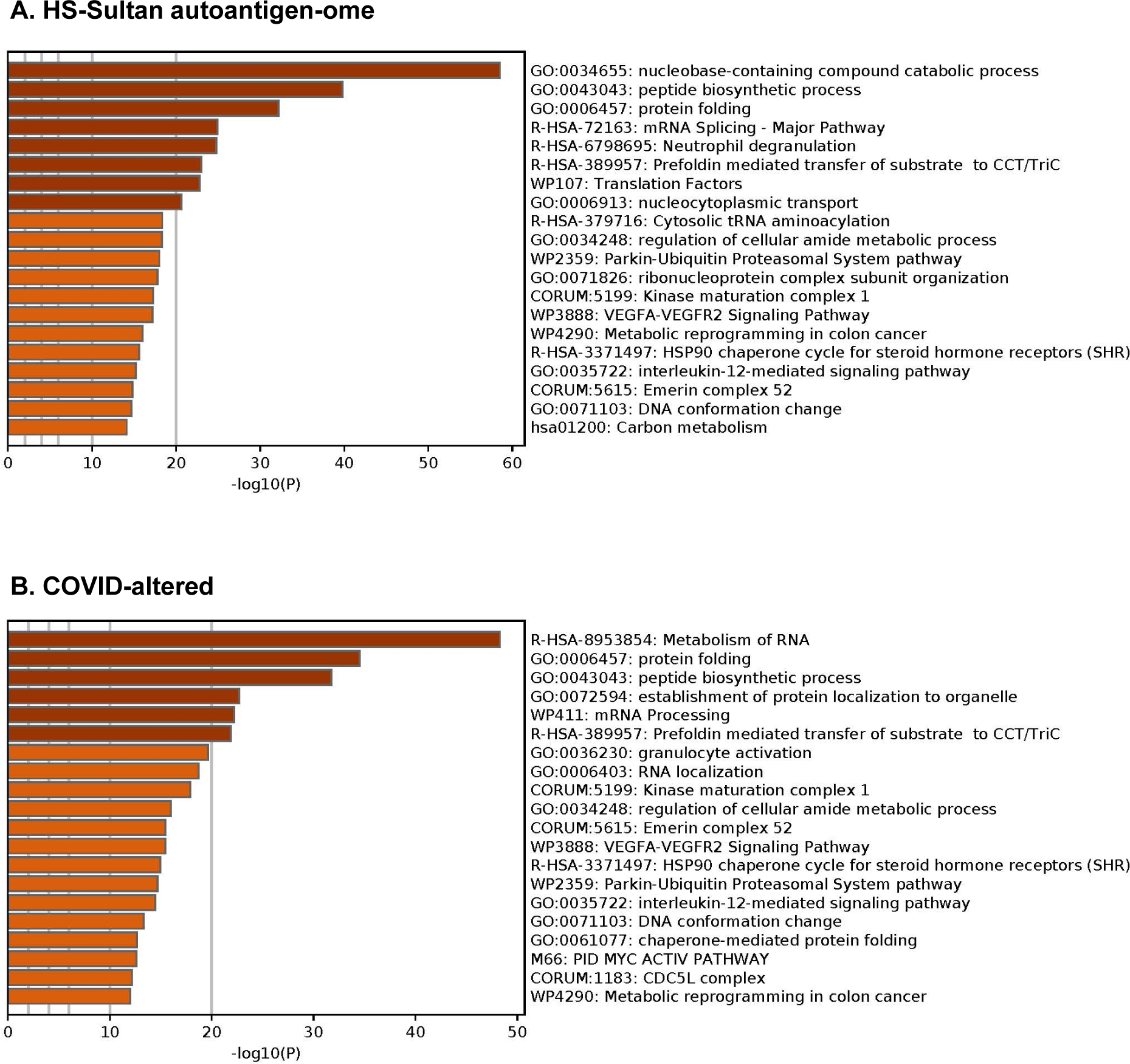
Top 20 enriched pathways and processes among COVID-altered autoAgs. Top: Pathways of 362 proteins identified by DS-affinity from HS-Sultan cells. Bottom: Pathways of 316 DS-affinity proteins that are altered in COVID.

The DS-affinity autoantigen-ome is dominated by several families of proteins. There are 24 proteasomal proteins, 22 spliceosome proteins, 14 hnRNPs, 13 aminoacyl-tRNA synthases (ligases), 13 translation initiation factors subunits, 12 ribosomal proteins, 10 heat shock proteins, 9 actin and actin-related proteins, 9 tubulins, 8 histones, 8 snRNPs, 7 T-complex proteins (CCT/TriC), 6 elongation factor subunits, and 6 14-3-3 proteins. The majority of the proteins in these families have been reported as autoAgs (Table 1). For example, all hnRNP and snRNP proteins identified by DS-affinity in this study are among the best-known nuclear autoAgs. Interestingly, autoAgs included in clinical diagnostic autoimmune disease ANA screening panels, such as SSB (lupus La), SNRPD1 (Sm D1), SNRPD3 (Sm D3), histones, and TOP1, are all identified in this study by DS-affinity enrichment from HS-Sultan cells.

In addition to proteins, such as ribosomal and ribonucleoproteins, that can be consistently identified from a variety of cell types, HS-Sultan B lymphoblast cells give rise to a large number of unique DS-affinity protein categories. In particular, many proteins associated with biomolecule biosynthesis are identified. Among them are proteins involved in inosine monophosphate and purine nucleotide biosynthesis (ATIC, GART, HPRT1, PAICS, PFAS, PPAT, SHMT1), amino acid biosynthesis (CS, IDH3A, PHGDH, PGAM1, PGAM2, PSPH), and carbohydrate biosynthesis and catabolic processes (ALDOA, ALDOC, ENO2, G6PD, GBE1, LDHA, TALDO1). There are also proteins involved in protein transport (ARF1, CSE1L, GDI1, GDI2, HMGB1, IPO5, KPNA2, RAB7A, RANBP1, RANBP6, SRP14, TNPO1, XPOT), dephosphorylation (NSF1C, PPP1R7, PPP2R1A, SET, SWAP70), and ubiquitination and de-ubiquitination (OTUB1, SHMT2, UBA1, UBA6, UBE2K, USP5). 17 of these 44 proteins are currently known autoAgs, while the remainder await further investigation. Overall, HS-Sultan cells appear to be especially rich in biosynthetic protein machinery, which may explain the rapid proliferation of these cells in Burkitt lymphoma.

Thirteen aminoacyl-tRNA synthetases were identified by DS-affinity from HS-Sultan cells, including AARS, DARS, ERPS, FARSB, GARS, HARS, KARS, NARS, PUS1, SARS, VARS, WARS, and YARS. Ten of these are already known autoAgs (Table 1), although we suspect that the remainder will also likely be autoAgs. This group of proteins are the culprits of antisynthetase syndrome, an autoimmune disease characterized by autoantibodies against one or multiple tRNA synthetases. Antisynthetase syndrome is a chronic disorder that can affect many parts of the body, with common symptoms including myositis, interstitial lung disease, polyarthritis, skin thickening and cracking of fingers and toes, or Raynaud disease. Antisynthetase syndrome has been reported in a case report of COVID-19 interstitial lung disease [39].

HS-Sultan cells are B lymphoblasts immortalized by Epstein-Barr virus (EBV) infection and carry viral DNA sequences. Using DS-affinity, we identified numerous proteins involved in DNA repair and the mitotic cell cycle, including CLTC, DCTN2, MCM2, MCM3, MCM4, MCM6, NSF1C, PNCA, PPAT, and SUGT1. Using DS-affinity, we also identified many proteins associated with telomerase maintenance, including TCP1, CCT2, CCT4, CCT5, CCT7, HNRNPA1, HNRPNA2B1, HNRNPC, HNRNPD, HNRNPU, HSP90AA1, HSP90AB1, PARP1, PTGES3, and XRCC5. Telomerase maintenance, which counteracts DNA damage response, cell cycle arrest, and apoptosis, is crucial for immortalization of cells with unlimited proliferative potential. Of these 25 proteins, 19 are known autoAgs, which indicates that proteins involved in telomerase maintenance, DNA repair, and cell cycle may be affected by EBV infection and become autoantigenic.

### DS-affinity autoantigen-ome related to SARS-CoV-2 infection

To find out whether DS-affinity autoAgs are affected in SARS-CoV-2 infection, we conducted similarity searches with currently available multi-omic COVID-19 data compiled in Coronascape (as of 02/22/2021) [17–38]. Among our 362 DS-affinity proteins, 315 (87.0%) are affected by SARS-CoV-2 infection (Table 1). Of these 315 proteins, 209 are up-regulated and 248 are down-regulated at protein and/or mRNA levels, and 95 are in the interactomes of individual SARS-CoV-2 viral proteins. Because the COVID-19 multi-omics data have been obtained with various methods from different infected cells or patients, there are proteins found up-regulated in some studies but down-regulated in other studies, but nevertheless, these proteins are affected by the infection and thus considered COVID-affected in our analysis (Supplemental Table 1). Of the 315 COVID-affected DS-affinity proteins, 186 (59.0%) are thus far confirmed autoAgs, while 129 are putative autoAgs (Table 1).

The COVID-affected DS-affinity proteins are highly connected (Fig. 3). By STRING analysis, these 315 proteins exhibit 2,507 interactions at high confidence level, which is significantly higher than randomly expected (1,002 interactions) with PPI enrichment p-value <1.0E-16. The proteins are primarily associated with RNA and mRNA processing, translation, vesicles, and vesicle-mediated transport (Fig. 3), which is consistent with our findings in other cell types [1, 2, 8]. In addition, these proteins are enriched in protein folding, peptide biosynthesis, granulocyte activation, emerin complex, IL-12 mediated signaling pathway, CDC5L complex, and metabolic reprogramming (Fig. 2B).

**Fig. 3.**
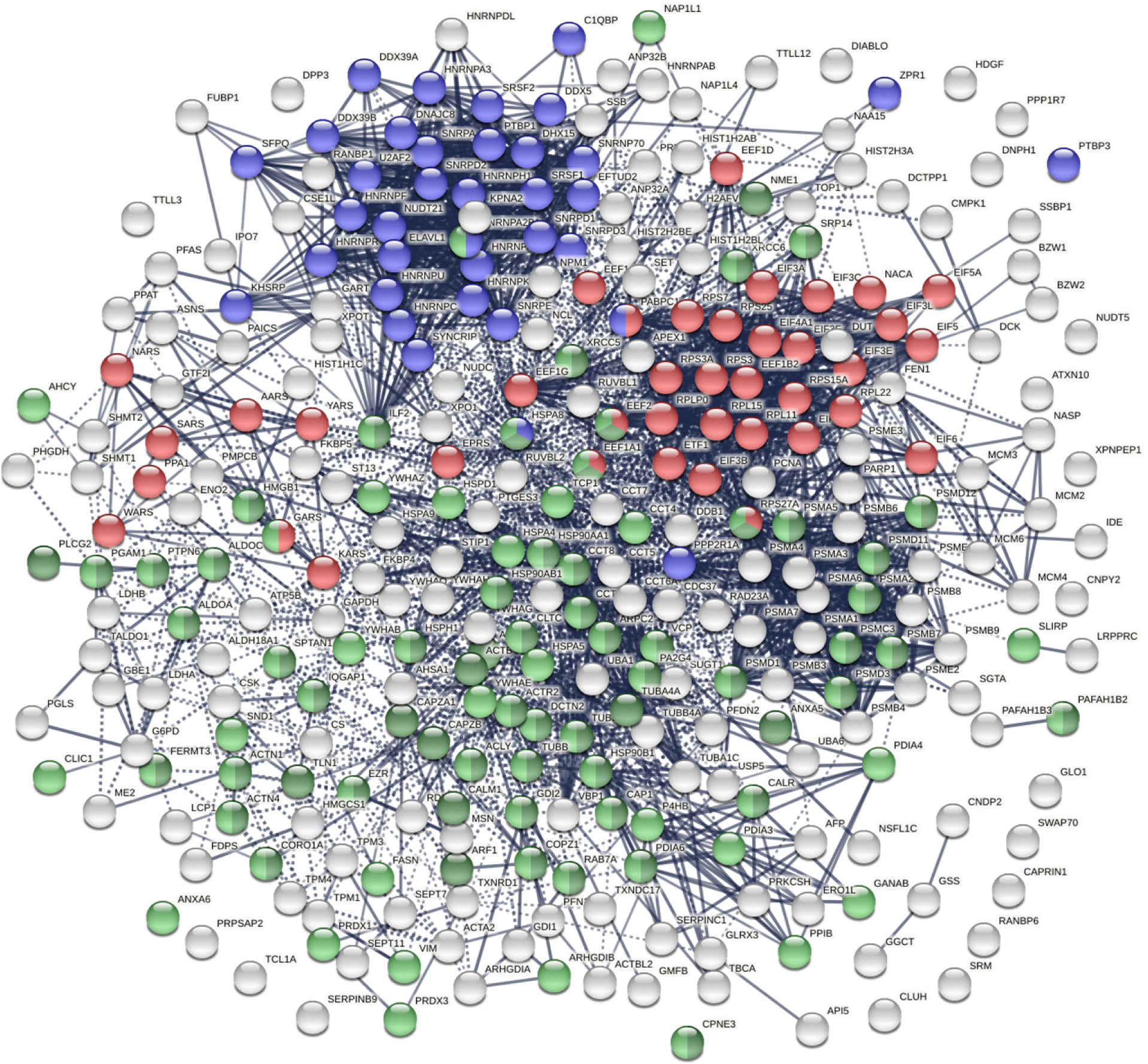
COVID-affected proteins shared with the HS-Sultan autoantigen-ome. Marked proteins are associated with RNA splicing (36 proteins, blue), translation (39 proteins, red), vesicle (77 proteins, green) and vesicle-mediated transport (62 proteins, dark green).

Twenty-one COVID-affected DS-affinity proteins are associated with mRNA splicing, including heterogenous nuclear ribonucleoproteins (hnRNP A1, A2B1, A3, AB, C, DL, F, H1, K, Q, R, and U), small nuclear ribonucleoproteins (SNRNP70, SNRPA, SNRPE, SNRPD1, SNRRPD2, SNRPD3), and splicing factors (SRSF1, SRSF2, SFPQ), all of which are well-known autoAgs.

Phosphorylation and ubiquitination changes induced by SARS-CoV-2 infection are posttranslational molecular alterations that may transform native proteins into potential autoAgs (Fig. 4), which is consistent with our previous findings [2]. Phosphorylation changes affected 80 COVID-altered DS-affinity proteins, including 8 hnRNPs, 4 initiation factors (EIF3A, 3B, 5), 3 elongation factors, 3 replication licensing factors (MCM2, 3, 4), SSB, XRCC6, and GTF2I. These phosphoproteins are associated with mRNA splicing, translation, telomere maintenance, DNA conformation change, and pre-replicative complex assembly. Ubiquitination changes affected 101 COVID-altered DS-affinity proteins, including 8 heat shock proteins, 5 initiation factors (EIF3E, 3F, 3I, 4A1, 5A), 4 CCT units, 4 14-3-3 proteins, 3 elongation factors, 3 histones, and 2 MCMS. These ubiquitinated proteins are associated with the nucleobase-containing compound catabolic process, RNA metabolism, cellular response to stress, prefoldin mediated transfer of substrate to CCT/TriC, and axon guidance.

**Fig. 4.**
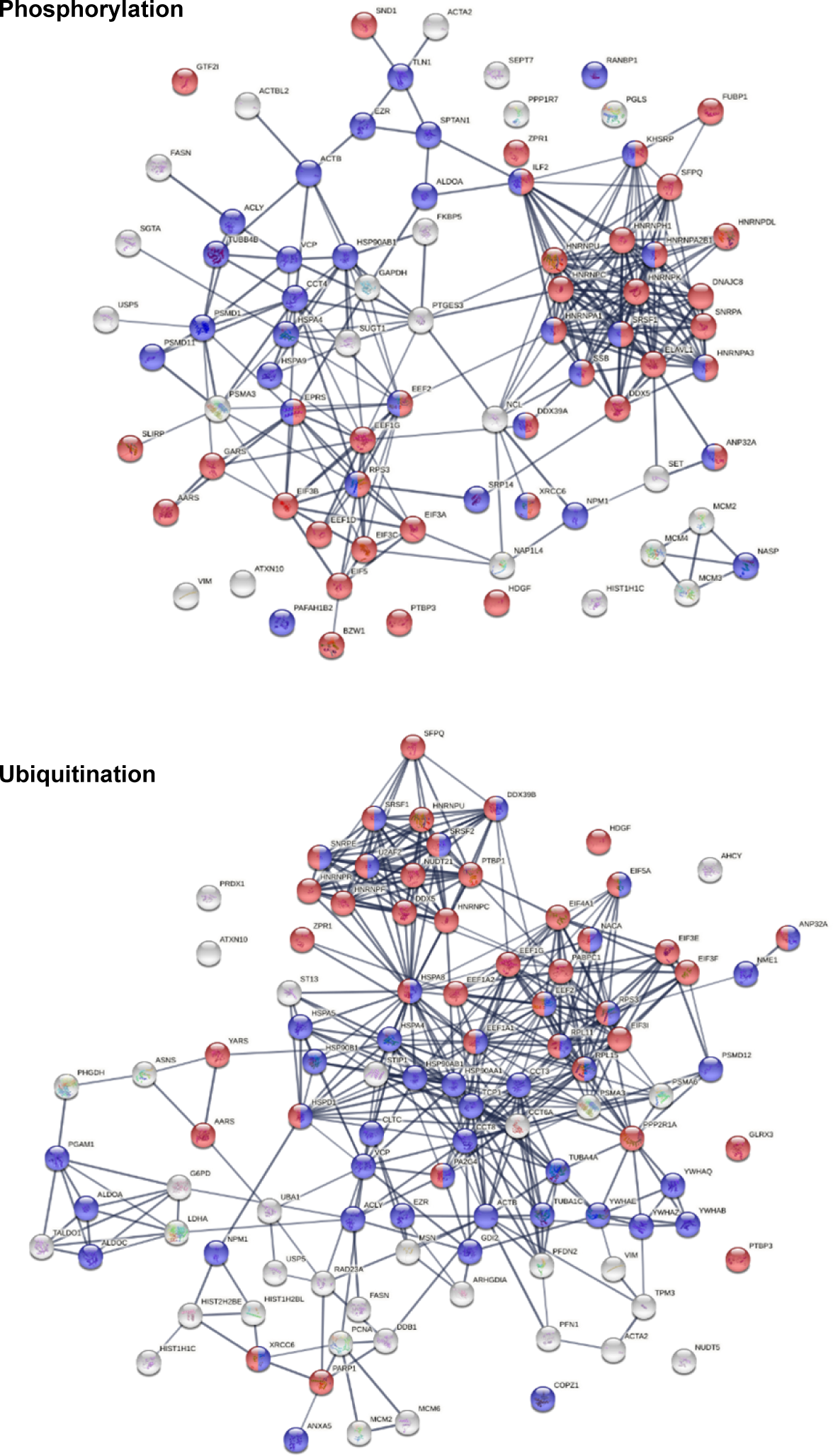
DS-affinity proteins that are altered by phosphorylation or ubiquitination in SARS- CoV-2 infection. Marked proteins are associated with gene expression (red) and transport (blue).

### AutoAgs that interact with SARS-CoV-2 components

There are 95 DS-affinity proteins found in the interactomes of various SARS-CoV-2 proteins (Fig. 5), meaning that these proteins can interact directly or indirectly with viral components. They appear to be intimately involved in protein metabolism, including 17 proteins related to peptide biosynthesis, 25 related to protein folding, 29 related to protein localization, and 22 related to proteolysis. In addition, these proteins are associated with the symbiont viral process, translational initiation, protein deubiquitination, protein stabilization, and posttranslational protein modification.

**Fig. 5.**
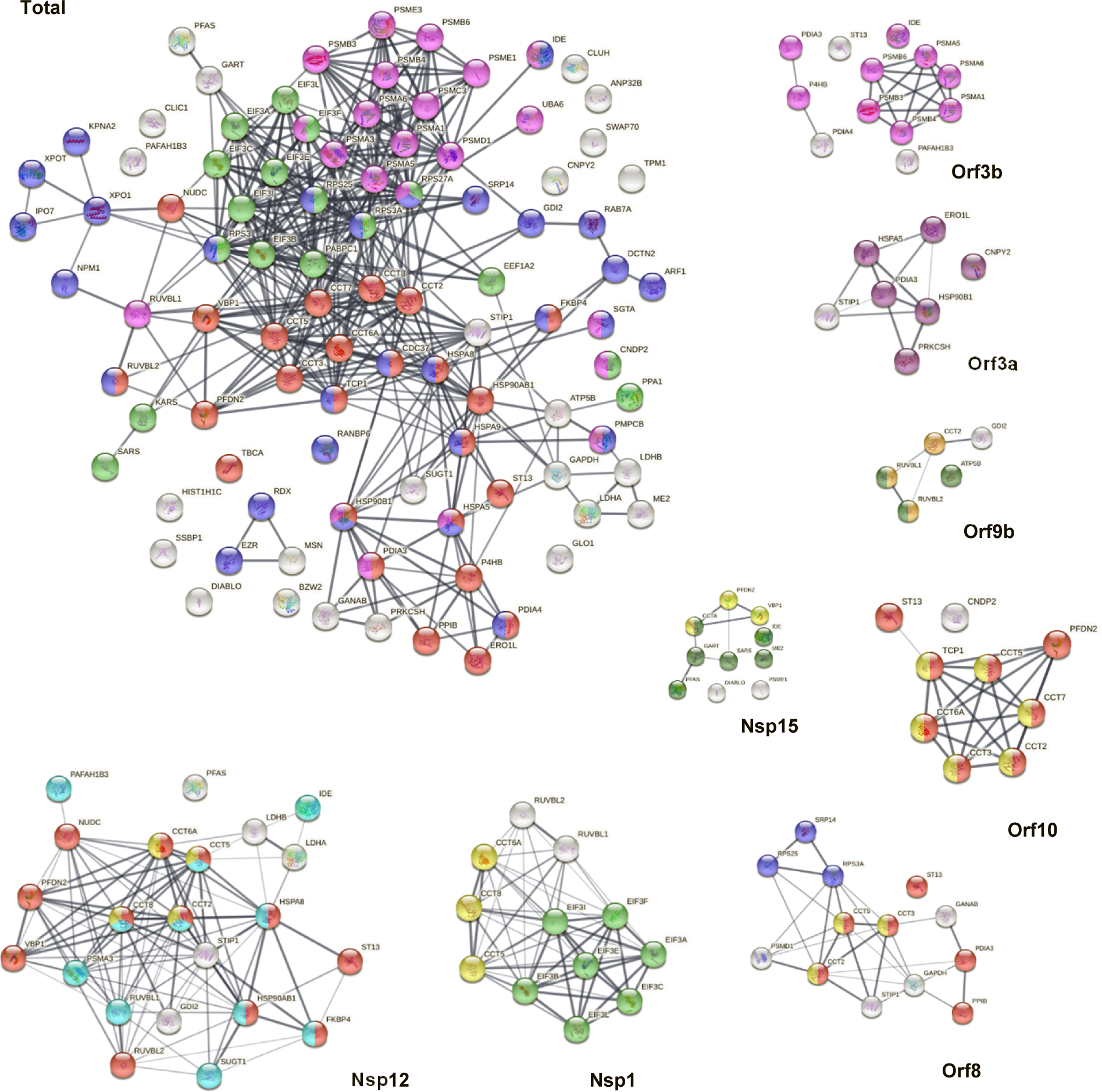
DS-affinity proteins in the SARS-CoV-2 interactomes. **Total:** marked proteins are involved in protein folding (25 proteins, red), peptide biosynthetic process (17 proteins, green), protein localization (29 proteins, blue), or proteolysis (22 proteins, pink). **Orf3b**: proteolysis (pink). **Orf3a**: endoplasmic reticulum (dark purple). **Orf9b**: nuclear function of prefoldin (amber), AAA+ ATPase domain or P-loop containing nucleoside triphosphate hydrolase (dark green). **Nsp15**: prefoldin-mediated transfer of substrate to CCT/TriC (yellow), nucleotide binding (dark green). **Orf10**: protein folding (red), CCT chaperonin (yellow). **Orf8:** protein folding (red), SRP-dependent cotranslational protein targeting to membrane (blue), CCT chaperonin (yellow). **Nsp1**: translation initiation (green), CCT chaperonin (yellow). **Nsp12**: protein folding (red), multi-organism process (aqua), CCT chaperonin (yellow).

The chaperonin-containing T-complex (CCT), also known as T-complex protein ring complex (TriC), is the chaperonin of eukaryotic cells. The human CCT/TriC complex is a two-ring barrel-like complex formed by 8 similar but distinct subunits. Remarkably, all 8 CCT subunits are identified by DS-affinity, and 7 of them are found in the SARS-CoV-2 interactomes: CCT2 (interacts with Nsp12, Orf8, Orf9b, Orf10), CCT5 (with Nsp1, Nsp12, Orf8, Orf10), CCT6A (with Nsp1, Nsp12, Orf10), CCT8 (with Nsp1, Nsp12, Nsp14), CCT3 (with Orf8, Orf10), TCP1 (with Orf10), and CCT7 (with Orf10). In total, 6 SARS-CoV-2 proteins interact with the host cell CCT chaperonin, with Orf10 interacting with 6 CCT subunits (Fig. 5). Furthermore, CCT subunits and PDFN2 (prefoldin subunit 2) are found together in the interactomes of Orf10, Nsp12, and Nsp15. CCTs assist the folding of proteins upon ATP hydrolysis, aiding in the folding of ∼10% of the proteome. PDFN2 binds the nascent polypeptide chain and promotes folding in an environment in which there are many competing pathways for nonnative proteins. Therefore, these findings suggest that SARS-CoV-2 exploits CCT complex and PDFN2 to ensure competitive folding of nonnative viral proteins in the host cells.

In addition to CCT/TriC, heat shock proteins (HSPs) are another group of chaperones frequently identified with DS-affinity. Ten HSPs are identified in this study, including HSPA4, HSPA5, HSPA8, HSPA9, HSPD1, HSPH1, HSP90AA1, HSP90AA2, HSP90AB1, and HSP9B1. All 10 are known autoAgs (Table 1). HSP8 interacts with Nsp2 and Nsp12. HSP90B1 (endoplasmin) interacts with Orf3a and Orf9c. HSPA9 interacts with N protein. HSP90AB1 interacts with Nsp12. Most strikingly, HSPA5 (GRP78, BiP) interacts with 9 SARS-CoV- 2 proteins, including S, E, M, Nsp2, Nsp4, Nsp6, Orf3a, Orf7a, and Orf7b. In addition, CDC37 (Hsp90 co-chaperone, hsp90 chaperone protein kinase-targeting) interacts with Nsp16. ST13 (Hsc70-interacting protein) interacts with 5 SARS-CoV-2 proteins (Nsp12, Orf3b, Orf6, Orf8, and Orf10). STIP1 (stress induced phosphoprotein 1, HSP90AA1 co-chaperone) interacts with 4 viral proteins (Nsp12, Orf3a, Orf8, and E).

The replication machinery of SARS-CoV-2 interacts with 41 different DS-affinity proteins. Nsp12, an RNA-dependent RNA polymerase and the central component of the replication machinery, interacts with the largest number (i.e., 22) of DS-affinity proteins (Fig. 5). Its cofactor Nsp7 interacts with 12 proteins and Nsp8 interacts with only one. The replication machine also includes a helicase (Nsp13), 2 ribonucleases (Nsp14 and Nsp15), 2 RNA-cap methyltransferases (Nsp14, Nsp16), and cofactor Nsp10. Nsp15 interacts with 10 DS-affinity proteins, Nsp16 interacts with 8 proteins, Nsp13 interacts with SRP14 and RDX, Nsp14 interacts with IDE and CCT8, and Nsp10 interacts with PSMA3. Nsp12-interacting DS-affinity proteins are strongly associated with protein folding, particularly prefoldin mediated transfer of substrates to CCT complex and cooperation of prefoldin and CCT in protein folding (Fig. 5). Nsp15-interacting proteins are also associated with prefoldin-mediated substrate transfer to CCT. DS-affinity proteins interacting with other individual viral replication components have no clear biological associations.

Orf3b of SARS-CoV-2 interacts with 12 DS-affinity proteins, including 6 proteasomal proteins, 3 protein disulfide-isomerases, IDE, ST13, and PAFAH1B3 (Fig. 5). Orf3a interacts with 7 proteins, including STIP1 (stress-induced-phosphoprotein 1) and 6 ER proteins (HSPA5, HSP90B1, CNPY2, ERO1L, PRKCSH, and PDIA3). CANPY2 prevents MIR-mediated MRLC ubiquitination and its subsequent proteasomal degradation. ERO1L (or ERO1A) is an oxidoreductase in disulfide bond formation in the ER. PRKCSH (glucosidase II subunit beta) cleaves sequentially the 2 innermost glucose residues from the Glc2Man9GlcNAc2 oligosaccharide precursor of immature glycoproteins. Based on the normal functions of their interacting proteins, Orf3a and Orf3b appear to affect host stress response and protein processing in the ER.

The S protein of SARS-CoV-2 is found to interact with HSPA5 (GRP78/BiP), PRKCSH, PRS27A (ubiquitin-40S ribosomal protein), MSN, and EZR. EZR and MSN are members of the ezrin-moesin-radixin (ERM) family, and its third member RDX is found to interact with Nsp13 of the virus. Moesin is localized to filopodia and other membranous protrusions that are important for cell-cell recognition, and ERM proteins connect the plasma membranes to the actin-based cytoskeleton. Actin and cytoskeleton proteins have been consistently found to be altered in SARS-CoV-2 infection in our previous studies [1, 2], and this finding suggests that ERM proteins facilitate the viral trafficking from host cell membrane to the cytoskeleton. All three ERM proteins are confirmed autoAgs.

Nsp1 is a major virulence factor of coronavirus. COVID-19 patients with autoantibodies are found to have higher levels of antibodies against SARS-CoV-2 Nsp1 protein [9]. Nsp1 has been reported to hijack the host 40S ribosome by inserting its C terminus into the mRNA entry tunnel, which effectively blocks RGI- dependent innate immune responses [40]. In this study, we found that Nsp1 interacts with 7 subunits of the translation initiation factor 3 complex (EIF3 A, B, C, E, F, I, L). EIF3 complex binds the 40S ribosome and serves as a scaffold for other initiation factors, auxiliary factors, and mRNA. Hence, our study extends previous reported activities of Nsp1 and shows that Nsp1 engages both the 40S ribosome and EIF3 to manipulate host protein translation.

A few interesting SARS-CoV-2-interacting DS-affinity proteins may provide clues to potential COVID-19 symptoms. PAFAH1B3 is a catalytic unit of the platelet-activating factor acetylhydrolase complex and plays important roles in platelet activation regulation and brain development, and it interacts with Nsp12, Nsp5, and Orf3b. Another subunit, PAFAH1B2, is altered in SARS-CoV-2 infection. Both this and our previous studies [2] have identified PAFAH1B2 and B3 as potential COVID-altered autoAgs, and their roles in COVID coagulopathy merit further investigation.

IDE (insulin degrading enzyme) is a ubiquitously expressed metalloprotease that degrades insulin, beta amyloid, and others. IDE interacts with 6 SARS-CoV-2 proteins (Nsp4, Nsp12, Nsp14, Nsp15, Nsp16, and Orf3b). Although its role in COVID remains unknown, IDE has been partially characterized in other viral infections. It is one of the host factors of hepatitis C virus [41], and it degrades HIV-1 p6 Gap protein and regulates virus replication in an Env-dependent manner [42]. In varicella zoster virus infection, the viral gE protein precursor associates with IDE, HSPA5, HSPA8, HSPD1, and PPIA in the ER of infected cells [43].

Interestingly, this group of ER proteins is also identified in this study, although we identified PPIB instead of PPIA. Although IDE has not yet formally been described as an autoAg, we have identified IDE in this and another study [2], and its importance for COVID-19 and autoimmunity merits further investigation.

### DS-affinity and B-cell-specific IgH-ER complex

Because HS-Sultan cells are derived from B lymphoblasts infected by Epstein-Barr virus, we compared the DS-affinity autoantigen-ome with single-cell mRNA expression profiles of B-cells from 7 patients hospitalized with COVID-19 [23]. We identified 39 DS-affinity proteins that are up-regulated at mRNA level in COVID B-cells, which include 7 heat shock proteins, 6 proteasomal proteins, 4 protein disulfide-isomerases, HDGF (heparin binding growth factor), CLIC1, CPNE3, SND1, TALDO1, TCL1A, and others (Fig. 6). These up-regulated proteins are primarily associated with protein processing in the ER and the proteasome. We also identified 21 DS-affinity proteins that are down-regulated in COVID B-cells, including 4 translation elongation factors, 2 translation initiation factors, 2 hnRNPs, 2 aminoacyl-tRNA synthetases, NACA, NAP1L1, and PABPC1. These down-regulated proteins are primarily associated with gene expression (Fig. 6).

**Fig. 6.**
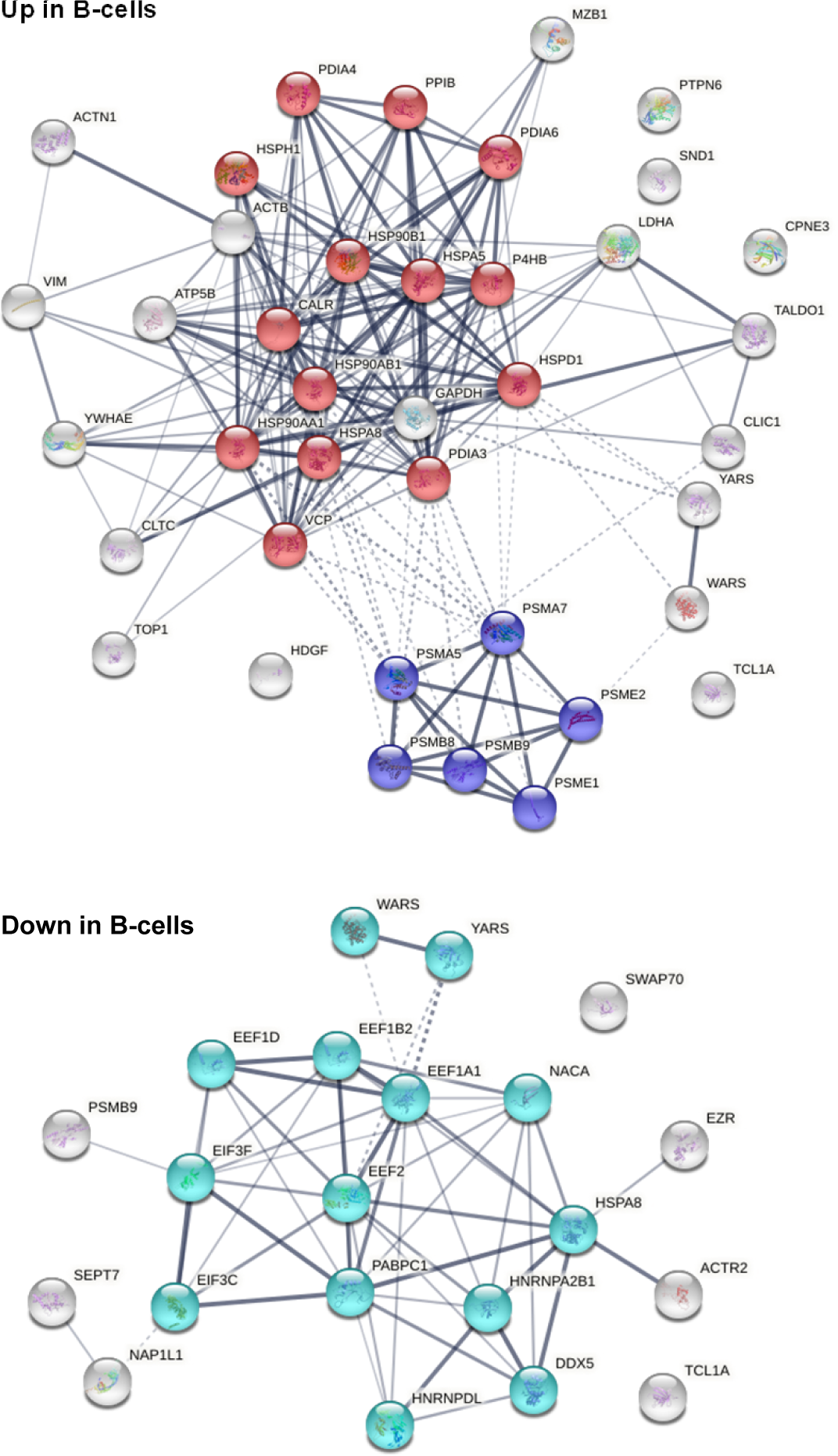
DS-affinity proteins from HS-Sultan cells that are up-regulated or down-regulated in B-cells of COVID-19 patients. **Top**: 39 up-regulated proteins. Red: protein processing in ER. Blue: proteasome. **Bottom**: 21 down-regulated proteins. Aqua: proteins involved in gene expression.

In particular, MZB1 (marginal zone B- and B1-cell-specific protein) is found up-regulated in B-cells from 5 of the 7 COVID-19 patients. MZB1 is a B-cell-specific ER-localized protein that is most abundantly expressed in marginal zone B-cells and B1-cells [44]. These cells are also termed innate-like B cells. They differ from follicular B-cells by their attenuated Ca^2+^ mobilization, fast antibody secretion, and increased cell adhesion. MZB1 plays important roles in humoral immunity and helps diversify peripheral B-cell functions by regulating calcium stores, antibody secretion, and integrin activation. MZB1 mRNA expression was found increased by >2-fold in B-cells of SLE patients with active disease [45]. High MZB1 gene expression predicted adverse prognosis in chronic lymphocytic leukemia, follicular lymphoma, and diffuse large B-cell lymphoma [46]. High prevalence of MZB1-positive plasma B-cells in tissue fibrosis was found in human lung and skin fibrosis, and MZB1 levels correlated positively with tissue IgG and negatively with diffusion capacity of the lung [47].

MZB1 is part of a B-cell-specific ER chaperone complex, associates with IgM heavy chain (IgH) and HSP90B1 (Grp94), and augments IgM assembly and secretion [48]. MZB1 is also found to augment the function of PDIA3 (ERp57) [44]. In this study, MZB1, HSP90B1, HSPA5 (Grp78, BiP), PDIA4, PDIA6, and CALR are found jointly up-regulated in the B-cells of the same 5 COVID-19 patients. This finding suggests that these 6 proteins are in the same IgH-associated ER complex. In addition, several other up-regulated ER proteins are identified, including HSP90AB1, HSPD1, HSPA8, PDIA3, P4HB, and PPIB.

Interestingly, in our study of murine pre-B1 lymphoblasts, we also found that DS interacts with the same IgH-associated multiprotein complex in the ER [5]. In addition, we had observed that DS interacts directly with GTF2I in murine pre-B1 cells, and GTF2I is also identified by DS-affinity in human B lymphoblast HS-Sultan cells in this study. GTF2I is a required gene transcription factor at the *IgH* gene locus. Pre-B1 cells, which express precursor B-cell receptors (preBCRs) that are polyreactive and autoreactive, are a critical check point in the development of mature autoreactive B cells. The Ig heavy chain (IgH) repertoire of autoantibodies is determined at the pre-B stage. Our previous findings from pre-B1 cells suggested that DS is a potential master regulator of IgH at both the gene and protein expression levels, i.e., DS recruits GTFI for *IgH* gene expression and engages IgH-associated ER complex for autoantibody production. The findings from this study provide further support for a key role of DS in regulating autoantibody production and autoreactive B1-cell development. Furthermore, the finding from B-cells of COVID-19 patients point to a potential significance of autoreactive B1 cells in COVID-induced autoimmunity.

## Conclusion

Exploiting the affinity between autoAgs and DS glycosaminoglycan, we identified 362 DS-affinity proteins from EBV-immortalized HS-Sultan cells. 201 of these DS-affinity proteins are already known autoAgs in a wide variety of autoimmune diseases and cancer. Of the 362, 315 DS-affinity proteins are affected by SARS-CoV-2 infection, and 186 COVID-affected DS-affinity proteins are known autoAgs. These COVID-altered proteins are largely affected by phosphorylation, ubiquitination, or interaction with viral protein components. They are associated with gene expression, mRNA processing, mRNA splicing, translation, protein folding, DNA replication fork, telomerase maintenance, chromosome organization, biosynthesis and catabolism of nucleobase-containing molecules and proteins, vesicles, and nucleocytoplasmic transport. CCT/TriC chaperonin, insulin degrading enzyme, and platelet-activating factor acetylhydrolase are found in the interactomes of multiple viral Nsp and Orf proteins. By integrating DS-affinity autoAgs with multi-omic data from COVID, our study suggests that viral infections can cause significant proteomic alterations, give rise to a diverse pool of autoAgs, and may lead to infection-induced autoimmune diseases. The COVID autoantigen-ome provided in this paper may serve as a molecular map and resource for investigating autoimmune phenomena of SARS-CoV-2 infection and its long-term sequelae. Understanding immunogenic proteins of COVID may also enhance vaccine target design.

## Materials and Methods

### HS-Sultan cell culture

The human B lymphoblast HS-Sultan cell line was obtained from the ATCC (Manassas, VA) and cultured in complete RPMI-1640 medium. The growth medium was supplemented with 10% fetal bovine serum and a penicillin-streptomycin-glutamine mixture (Thermo Fisher). The cells were grown at 37 °C in a CO_2_ incubator, and about 300 million cells were harvested for the study.

### Protein extraction

Protein extraction was performed as previously described [4]. In brief, HS-Sultan cells were lysed with 50 mM phosphate buffer (pH 7.4) containing the Roche Complete Mini protease inhibitor cocktail and then homogenized on ice with a microprobe sonicator until the turbid mixture turned nearly clear with no visible cells left. The homogenate was centrifuged at 10,000 g at 4 °C for 20 min, and the total protein extract in the supernatant was collected. Protein concentration was measured by absorbance at 280 nm using a NanoDrop UV-Vis spectrometer (Thermo Fisher).

### DS-Sepharose resin preparation

The DS-affinity resins were synthesized as previously described [4, 8]. In brief, 20 ml of EAH Sepharose 4B resins (GE Healthcare Life Sciences) were washed with distilled water three times and mixed with 100 mg of DS (Sigma-Aldrich) in 10 ml of 0.1 M MES buffer, pH 5.0. About 100 mg of N-(3-dimethylaminopropyl)- N’-ethylcarbodiimide hydrochloride (Sigma-Aldrich) powder was added, and another 100 mg was added after 8 h of reaction. The reaction proceeded by mixing on a rocker at 25 °C for 16 h. The coupled resins were washed with water and equilibrated with 0.5 M NaCl in 0.1 M acetate (pH 5.0) and 0.5 M NaCl in 0.1 M Tris (pH 8.0).

### DS-affinity fractionation

The total proteomes extracted from HS-Sultan cells were fractionated in a DS-Sepharose column in a manner similar to previously described [4]. About 40 mg of proteins in 40 ml of 10 mM phosphate buffer (pH 7.4; buffer A) were loaded onto the DS-affinity column at a rate of 1 ml/min. Unbound and weakly bound proteins were removed with 60 ml of buffer A and then 40 ml of 0.2 M NaCl in buffer A. The remaining bound proteins were eluted in steps with 40 ml 0.5 M NaCl and then with 40 ml 1.0 M NaCl in buffer A. Fractions were desalted and concentrated with 5-kDa cut-off Vivaspin centrifugal filters (Sartorius). Fractionated proteins were separated in 1-D SDS-PAGE in 4-12% Bis-Tris gels, and the gel lanes were divided into two or three sections for mass spectrometric sequencing.

### Mass spectrometry sequencing

Protein sequencing was performed at the Taplin Biological Mass Spectrometry Facility at Harvard Medical School. Proteins in gels were digested with sequencing-grade trypsin (Promega) at 4 °C for 45 min. Tryptic peptides were separated in a nano-scale C18 HPLC capillary column and analyzed in an LTQ linear ion-trap mass spectrometer (Thermo Fisher). Peptide sequences and protein identities were assigned by matching the measured fragmentation pattern with proteins or translated nucleotide databases using Sequest. All data were manually inspected. Proteins with ≥2 peptide matches were considered positively identified.

### COVID data comparison

DS-affinity proteins were compared with currently available COVID-19 multi-omic data compiled in the Coronascape database (as of 02/22/2021) [17–38]. These data have been obtained with proteomics, phosphoproteomics, interactome, ubiquitome, and RNA-seq techniques. Up- and down-regulated proteins or genes were identified by comparing cells infected vs. uninfected by SARS-CoV-2 or COVID-19 patients vs. healthy controls. Similarity searches were conducted to identify DS-affinity proteins that are up- and/or down-regulated in viral infection at any omic level.

### Protein network analysis

Protein-protein interactions were analyzed by STRING [49]. Interactions include both direct physical interaction and indirect functional associations, which are derived from genomic context predictions, high-throughput lab experiments, co-expression, automated text mining, and previous knowledge in databases. Each interaction is annotated with a confidence score from 0 to 1, with 1 being the highest, indicating the likelihood of an interaction to be true. Pathways and processes enrichment were analyzed with Metascape [17], which utilize various ontology sources such as KEGG Pathway, GO Biological Process, Reactome Gene Sets, Canonical Pathways, CORUM, TRRUST, and DiGenBase. All genes in the genome were used as the enrichment background. Terms with a p-value <0.01, a minimum count of 3, and an enrichment factor (ratio between the observed counts and the counts expected by chance) >1.5 were collected and grouped into clusters based on their membership similarities. The most statistically significant term within a cluster was chosen to represent the cluster.

### Autoantigen literature text mining

Every DS-affinity protein identified in this study was searched for specific autoantibodies reported in the PubMed literature. Search keywords included the MeSH keyword “autoantibodies”, the protein name or its gene symbol, or alternative names and symbols. Only proteins for which specific autoantibodies are reported in PubMed-listed journal articles were considered “confirmed” autoAgs in this study.

## Acknowledgements

We thank Jung-hyun Rho for technical assistance with experiments. We thank Ross Tomaino and the Taplin Biological Mass Spectrometry facility of Harvard Medical School for expert service with protein sequencing.

## Competing interest statement

JYW is the founder and Chief Scientific Officer of Curandis. WZ was supported by the NIH and declares no competing interests. MWR and VBR are volunteers of Curandis. MHR is a member of the Scientific Advisory Boards of Trans-Hit, Proscia, and Universal DX, but these companies have no relation to the study.

### Authors’ contributions

JYW directed the study and wrote the manuscript. WZ performed part of the experiments. MWR and VBR assisted with data analysis and manuscript preparation. MHR consulted on the study and edited the manuscript. All authors have approved the manuscript.

## References

1. Wang JY, Zhang W, Roehrl MW, Roehrl VB, Roehrl MH. An Autoantigen Atlas from Human Lung HFL1 Cells Offers Clues to Neurological and Diverse Autoimmune Manifestations of COVID-19. bioRxiv. 2021:2021.01.24.427965. doi: 10.1101/2021.01.24.427965.

2. Wang JY, Zhang W, Roehrl MW, Roehrl VB, Roehrl MH. An Autoantigen Profile of Human A549 Lung Cells Reveals Viral and Host Etiologic Molecular Attributes of Autoimmunity in COVID-19. bioRxiv. 2021:2021.02.21.432171. doi: 10.1101/2021.02.21.432171.

3. Wang JY, Lee J, Yan M, Rho JH, Roehrl MH. Dermatan sulfate interacts with dead cells and regulates CD5(+) B-cell fate: implications for a key role in autoimmunity. Am J Pathol. 2011;178(5):2168–76. doi: 10.1016/j.ajpath.2011.01.028. PubMed PMID: 21514431; PubMed Central PMCID: PMCPMC3081202.

4. Rho JH, Zhang W, Murali M, Roehrl MH, Wang JY. Human proteins with affinity for dermatan sulfate have the propensity to become autoantigens. Am J Pathol. 2011;178(5):2177–90. doi: 10.1016/j.ajpath.2011.01.031. PubMed PMID: 21514432; PubMed Central PMCID: PMCPMC3081203.

5. Lee J, Rho J-h, Roehrl MH, Wang JY. Dermatan Sulfate Is a Potential Master Regulator of IgH via Interactions with Pre-BCR, GTF2I, and BiP ER Complex in Pre-B Lymphoblasts. bioRxiv. 2021:2021.01.18.427153. doi: 10.1101/2021.01.18.427153.

6. Zhang W, Rho JH, Roehrl MH, Wang JY. A comprehensive autoantigen-ome of autoimmune liver diseases identified from dermatan sulfate affinity enrichment of liver tissue proteins. BMC immunology. 2019;20(1):21. doi: 10.1186/s12865-019-0304-1. PubMed PMID: 31242852; PubMed Central PMCID: PMCPMC6595630.

7. Zhang W, Rho JH, Roehrl MW, Roehrl MH, Wang JY. A repertoire of 124 potential autoantigens for autoimmune kidney diseases identified by dermatan sulfate affinity enrichment of kidney tissue proteins. PloS one. 2019;14(6):e0219018. doi: 10.1371/journal.pone.0219018. PubMed PMID: 31237920; PubMed Central PMCID: PMCPMC6592568.

8. Wang JY, Zhang W, Rho JH, Roehrl MW, Roehrl MH. A proteomic repertoire of autoantigens identified from the classic autoantibody clinical test substrate HEp-2 cells. Clinical proteomics. 2020;17:35. Epub 2020/09/26. doi: 10.1186/s12014-020-09298-3. PubMed PMID: 32973414; PubMed Central PMCID: PMCPMC7507713.

9. Chang SE, Feng A, Meng W, Apostolidis SA, Mack E, Artandi M, et al. New-Onset IgG Autoantibodies in Hospitalized Patients with COVID-19. medRxiv : the preprint server for health sciences. 2021. Epub 2021/02/04. doi: 10.1101/2021.01.27.21250559. PubMed PMID: 33532787; PubMed Central PMCID: PMCPMC7852238.

10. Bastard P, Rosen LB, Zhang Q, Michailidis E, Hoffmann HH, Zhang Y, et al. Autoantibodies against type I IFNs in patients with life-threatening COVID-19. Science (New York, NY). 2020;370(6515). Epub 2020/09/26. doi:10.1126/science.abd4585. PubMed PMID: 32972996.

11. Consiglio CR, Cotugno N, Sardh F, Pou C, Amodio D, Rodriguez L, et al. The Immunology of Multisystem Inflammatory Syndrome in Children with COVID-19. Cell. 2020;183(4):968–81.e7. Epub 2020/09/24. doi: 10.1016/j.cell.2020.09.016. PubMed PMID: 32966765; PubMed Central PMCID: PMCPMC7474869.

12. Gruber CN, Patel RS, Trachtman R, Lepow L, Amanat F, Krammer F, et al. Mapping Systemic Inflammation and Antibody Responses in Multisystem Inflammatory Syndrome in Children (MIS-C). Cell. 2020;183(4):982–95.e14. Epub 2020/09/30. doi: 10.1016/j.cell.2020.09.034. PubMed PMID: 32991843; PubMed Central PMCID: PMCPMC7489877.

13. Franke C, Ferse C, Kreye J, Reincke SM, Sanchez-Sendin E, Rocco A, et al. High frequency of cerebrospinal fluid autoantibodies in COVID-19 patients with neurological symptoms. Brain, behavior, and immunity. 2020. Epub 2020/12/29. doi: 10.1016/j.bbi.2020.12.022. PubMed PMID: 33359380; PubMed Central PMCID: PMCPMC7834471.

14. Zhou Y, Han T, Chen J, Hou C, Hua L, He S, et al. Clinical and Autoimmune Characteristics of Severe and Critical Cases of COVID-19. Clinical and translational science. 2020;13(6):1077–86. Epub 2020/04/22. doi: 10.1111/cts.12805. PubMed PMID: 32315487; PubMed Central PMCID: PMCPMC7264560.

15. Sacchi MC, Tamiazzo S, Stobbione P, Agatea L, De Gaspari P, Stecca A, et al. SARS-CoV-2 Infection as a trigger of autoimmune response. Clinical and translational science. 2020. Epub 2020/12/12. doi: 10.1111/cts.12953. PubMed PMID: 33306235.

16. Lerma LA, Chaudhary A, Bryan A, Morishima C, Wener MH, Fink SL. Prevalence of autoantibody responses in acute coronavirus disease 2019 (COVID-19). J Transl Autoimmun. 2020;3:100073. Epub 2020/12/03. doi: 10.1016/j.jtauto.2020.100073. PubMed PMID: 33263103; PubMed Central PMCID: PMCPMC7691817.

17. Zhou Y, Zhou B, Pache L, Chang M, Khodabakhshi AH, Tanaseichuk O, et al. Metascape provides a biologist-oriented resource for the analysis of systems-level datasets. Nat Commun. 2019;10(1):1523. doi: 10.1038/s41467-019-09234-6. PubMed PMID: 30944313; PubMed Central PMCID: PMCPMC6447622.

18. Zhang JY, Wang XM, Xing X, Xu Z, Zhang C, Song JW, et al. Single-cell landscape of immunological responses in patients with COVID-19. Nat Immunol. 2020;21(9):1107–18. Epub 2020/08/14. doi: 10.1038/s41590-020-0762-x. PubMed PMID: 32788748.

19. Davies JP, Almasy KM, McDonald EF, Plate L. Comparative Multiplexed Interactomics of SARS-CoV- 2 and Homologous Coronavirus Nonstructural Proteins Identifies Unique and Shared Host-Cell Dependencies. ACS infectious diseases. 2020;6(12):3174–89. Epub 2020/12/03. doi: 10.1021/acsinfecdis.0c00500. PubMed PMID: 33263384; PubMed Central PMCID: PMCPMC7724760.

20. Klann K, Bojkova D, Tascher G, Ciesek S, Münch C, Cinatl J. Growth Factor Receptor Signaling Inhibition Prevents SARS-CoV-2 Replication. Molecular cell. 2020;80(1):164–74.e4. Epub 2020/09/03. doi: 10.1016/j.molcel.2020.08.006. PubMed PMID: 32877642; PubMed Central PMCID: PMCPMC7418786.

21. Sun J, Ye F, Wu A, Yang R, Pan M, Sheng J, et al. Comparative Transcriptome Analysis Reveals the Intensive Early Stage Responses of Host Cells to SARS-CoV-2 Infection. Frontiers in microbiology. 2020;11:593857. Epub 2020/12/17. doi: 10.3389/fmicb.2020.593857. PubMed PMID: 33324374; PubMed Central PMCID: PMCPMC7723856.

22. Bojkova D, Klann K, Koch B, Widera M, Krause D, Ciesek S, et al. Proteomics of SARS-CoV-2- infected host cells reveals therapy targets. Nature. 2020;583(7816):469-72. Epub 2020/05/15. doi: 10.1038/s41586-020-2332-7. PubMed PMID: 32408336.

23. Wilk AJ, Rustagi A, Zhao NQ, Roque J, Martínez-Colón GJ, McKechnie JL, et al. A single-cell atlas of the peripheral immune response in patients with severe COVID-19. Nature medicine. 2020;26(7):1070–6. Epub 2020/06/10. doi: 10.1038/s41591-020-0944-y. PubMed PMID: 32514174; PubMed Central PMCID: PMCPMC7382903.

24. Lieberman NAP, Peddu V, Xie H, Shrestha L, Huang ML, Mears MC, et al. In vivo antiviral host transcriptional response to SARS-CoV-2 by viral load, sex, and age. PLoS biology. 2020;18(9):e3000849. Epub 2020/09/09. doi: 10.1371/journal.pbio.3000849. PubMed PMID: 32898168; PubMed Central PMCID: PMCPMC7478592.

25. Riva L, Yuan S, Yin X, Martin-Sancho L, Matsunaga N, Pache L, et al. Discovery of SARS-CoV-2 antiviral drugs through large-scale compound repurposing. Nature. 2020;586(7827):113-9. Epub 2020/07/25. doi: 10.1038/s41586-020-2577-1. PubMed PMID: 32707573; PubMed Central PMCID: PMCPMC7603405.

26. Bouhaddou M, Memon D, Meyer B, White KM, Rezelj VV, Correa Marrero M, et al. The Global Phosphorylation Landscape of SARS-CoV-2 Infection. Cell. 2020;182(3):685–712.e19. Epub 2020/07/10. doi: 10.1016/j.cell.2020.06.034. PubMed PMID: 32645325; PubMed Central PMCID: PMCPMC7321036.

27. Blanco-Melo D, Nilsson-Payant BE, Liu WC, Uhl S, Hoagland D, Møller R, et al. Imbalanced Host Response to SARS-CoV-2 Drives Development of COVID-19. Cell. 2020;181(5):1036–45.e9. Epub 2020/05/18. doi: 10.1016/j.cell.2020.04.026. PubMed PMID: 32416070; PubMed Central PMCID: PMCPMC7227586.

28. Shen B, Yi X, Sun Y, Bi X, Du J, Zhang C, et al. Proteomic and Metabolomic Characterization of COVID-19 Patient Sera. Cell. 2020;182(1):59–72.e15. Epub 2020/06/04. doi: 10.1016/j.cell.2020.05.032. PubMed PMID: 32492406; PubMed Central PMCID: PMCPMC7254001.

29. Lamers MM, Beumer J, van der Vaart J, Knoops K, Puschhof J, Breugem TI, et al. SARS-CoV-2 productively infects human gut enterocytes. Science (New York, NY). 2020;369(6499):50-4. Epub 2020/05/03. doi: 10.1126/science.abc1669. PubMed PMID: 32358202; PubMed Central PMCID: PMCPMC7199907.

30. Gordon DE, Jang GM, Bouhaddou M, Xu J, Obernier K, White KM, et al. A SARS-CoV-2 protein interaction map reveals targets for drug repurposing. Nature. 2020;583(7816):459-68. Epub 2020/05/01. doi: 10.1038/s41586-020-2286-9. PubMed PMID: 32353859; PubMed Central PMCID: PMCPMC7431030.

31. Xiong Y, Liu Y, Cao L, Wang D, Guo M, Jiang A, et al. Transcriptomic characteristics of bronchoalveolar lavage fluid and peripheral blood mononuclear cells in COVID-19 patients. Emerging microbes & infections. 2020;9(1):761–70. Epub 2020/04/02. doi: 10.1080/22221751.2020.1747363. PubMed PMID: 32228226; PubMed Central PMCID: PMCPMC7170362.

32. Vanderheiden A, Ralfs P, Chirkova T, Upadhyay AA, Zimmerman MG, Bedoya S, et al. Type I and Type III Interferons Restrict SARS-CoV-2 Infection of Human Airway Epithelial Cultures. Journal of virology. 2020;94(19). Epub 2020/07/24. doi: 10.1128/jvi.00985-20. PubMed PMID: 32699094; PubMed Central PMCID: PMCPMC7495371.

33. Appelberg S, Gupta S, Svensson Akusjärvi S, Ambikan AT, Mikaeloff F, Saccon E, et al. Dysregulation in Akt/mTOR/HIF-1 signaling identified by proteo-transcriptomics of SARS-CoV-2 infected cells. Emerging microbes & infections. 2020;9(1):1748–60. Epub 2020/07/22. doi: 10.1080/22221751.2020.1799723. PubMed PMID: 32691695; PubMed Central PMCID: PMCPMC7473213.

34. Stukalov A, Girault V, Grass V, Bergant V, Karayel O, Urban C, et al. Multi-level proteomics reveals host-perturbation strategies of SARS-CoV-2 and SARS-CoV. bioRxiv : the preprint server for biology. 2020:2020.06.17.156455. doi: 10.1101/2020.06.17.156455.

35. Emanuel W, Kirstin M, Vedran F, Asija D, Theresa GL, Roberto A, et al. Bulk and single-cell gene expression profiling of SARS-CoV-2 infected human cell lines identifies molecular targets for therapeutic intervention. bioRxiv. 2020:2020.05.05.079194. doi: 10.1101/2020.05.05.079194.

36. Li Y, Wang Y, Liu H, Sun W, Ding B, Zhao Y, et al. Urine Proteome of COVID-19 Patients. medRxiv. 2020:2020.05.02.20088666. doi: 10.1101/2020.05.02.20088666.

37. Liao M, Liu Y, Yuan J, Wen Y, Xu G, Zhao J, et al. Single-cell landscape of bronchoalveolar immune cells in patients with COVID-19. Nature medicine. 2020;26(6):842–4. Epub 2020/05/14. doi: 10.1038/s41591-020-0901-9. PubMed PMID: 32398875.

38. Laurent EMN, Sofianatos Y, Komarova A, Gimeno J-P, Tehrani PS, Kim D-K, et al. Global BioID-based SARS-CoV-2 proteins proximal interactome unveils novel ties between viral polypeptides and host factors involved in multiple COVID19-associated mechanisms. bioRxiv. 2020:2020.08.28.272955. doi: 10.1101/2020.08.28.272955.

39. Alfraji N, Mazahir U, Chaudhri M, Miskoff J. Anti-synthetase syndrome: a rare and challenging diagnosis for bilateral ground-glass opacities-a case report with literature review. BMC pulmonary medicine. 2021;21(1):11. Epub 2021/01/08. doi: 10.1186/s12890-020-01388-0. PubMed PMID: 33407281; PubMed Central PMCID: PMCPMC7787399.

40. Thoms M, Buschauer R, Ameismeier M, Koepke L, Denk T, Hirschenberger M, et al. Structural basis for translational shutdown and immune evasion by the Nsp1 protein of SARS-CoV-2. Science (New York, NY). 2020;369(6508):1249–55. Epub 2020/07/19. doi: 10.1126/science.abc8665. PubMed PMID: 32680882; PubMed Central PMCID: PMCPMC7402621.

41. Li Q, Zhang YY, Chiu S, Hu Z, Lan KH, Cha H, et al. Integrative functional genomics of hepatitis C virus infection identifies host dependencies in complete viral replication cycle. PLoS pathogens. 2014;10(5):e1004163. Epub 2014/05/24. doi: 10.1371/journal.ppat.1004163. PubMed PMID: 24852294; PubMed Central PMCID: PMCPMC4095987.

42. Hahn F, Schmalen A, Setz C, Friedrich M, Schlößer S, Kölle J, et al. Proteolysis of mature HIV-1 p6 Gag protein by the insulin-degrading enzyme (IDE) regulates virus replication in an Env-dependent manner. PloS one. 2017;12(4):e0174254. Epub 2017/04/08. doi: 10.1371/journal.pone.0174254. PubMed PMID: 28388673; PubMed Central PMCID: PMCPMC5384750.

43. Carpenter JE, Jackson W, de Souza GA, Haarr L, Grose C. Insulin-degrading enzyme binds to the nonglycosylated precursor of varicella-zoster virus gE protein found in the endoplasmic reticulum. Journal of virology. 2010;84(2):847–55. Epub 2009/10/30. doi: 10.1128/jvi.01801-09. PubMed PMID: 19864391; PubMed Central PMCID: PMCPMC2798375.

44. Flach H, Rosenbaum M, Duchniewicz M, Kim S, Zhang SL, Cahalan MD, et al. Mzb1 protein regulates calcium homeostasis, antibody secretion, and integrin activation in innate-like B cells. Immunity. 2010;33(5):723–35. Epub 2010/11/26. doi: 10.1016/j.immuni.2010.11.013. PubMed PMID: 21093319; PubMed Central PMCID: PMCPMC3125521.

45. Miyagawa-Hayashino A, Yoshifuji H, Kitagori K, Ito S, Oku T, Hirayama Y, et al. Increase of MZB1 in B cells in systemic lupus erythematosus: proteomic analysis of biopsied lymph nodes. Arthritis research & therapy. 2018;20(1):13. Epub 2018/02/01. doi: 10.1186/s13075-018-1511-5. PubMed PMID: 29382365; PubMed Central PMCID: PMCPMC5791339.

46. Herold T, Mulaw MA, Jurinovic V, Seiler T, Metzeler KH, Dufour A, et al. High expression of MZB1 predicts adverse prognosis in chronic lymphocytic leukemia, follicular lymphoma and diffuse large B-cell lymphoma and is associated with a unique gene expression signature. Leukemia & lymphoma. 2013;54(8):1652–7. Epub 2012/11/30. doi: 10.3109/10428194.2012.753445. PubMed PMID: 23189934.

47. Schiller HB, Mayr CH, Leuschner G, Strunz M, Staab-Weijnitz C, Preisendörfer S, et al. Deep Proteome Profiling Reveals Common Prevalence of MZB1-Positive Plasma B Cells in Human Lung and Skin Fibrosis. American journal of respiratory and critical care medicine. 2017;196(10):1298–310. Epub 2017/06/28. doi: 10.1164/rccm.201611-2263OC. PubMed PMID: 28654764; PubMed Central PMCID: PMCPMC6913086.

48. Rosenbaum M, Andreani V, Kapoor T, Herp S, Flach H, Duchniewicz M, et al. MZB1 is a GRP94 cochaperone that enables proper immunoglobulin heavy chain biosynthesis upon ER stress. Genes & development. 2014;28(11):1165–78. Epub 2014/06/04. doi: 10.1101/gad.240762.114. PubMed PMID: 24888588; PubMed Central PMCID: PMCPMC4052763.

49. Szklarczyk D, Gable AL, Lyon D, Junge A, Wyder S, Huerta-Cepas J, et al. STRING v11: protein-protein association networks with increased coverage, supporting functional discovery in genome-wide experimental datasets. Nucleic Acids Res. 2019;47(D1):D607–D13. doi: 10.1093/nar/gky1131. PubMed PMID: 30476243; PubMed Central PMCID: PMCPMC6323986.

